# A multi-omic *Nicotiana benthamiana* resource for fundamental research and biotechnology

**DOI:** 10.1101/2022.12.30.521993

**Authors:** Buddhini Ranawaka, Jiyuan An, Michał T. Lorenc, Hyungtaek Jung, Maria Sulli, Giuseppe Aprea, Sally Roden, Satomi Hayashi, Tal Cooper, Zacharie LeBlanc, Victor Llaca, Diego Orzaez, Aureliano Bombarely, Julia Bally, Christopher Winefield, Giovanni Giuliano, Peter M. Waterhouse

## Abstract

*Nicotiana benthamiana* is an invaluable model plant and biotechnology platform. To further improve its usefulness and versatility, we have produced high quality chromosome level genome assemblies and multi-omic datasets for both the ubiquitously used LAB strain and a distantly related wild accession, QLD, as well as mapping their single nucleotide polymorphisms with two additional laboratory and four additional wild accessions. LAB and QLD have many genetic, functional, and metabolic differences. These coupled with their high inter-fertility and equally efficient transient and stable transformation and genome editing provide a powerful resource partnership. Their ∼3Gb allotetraploid genomes show advanced diploidisation with major chromosome loss and rearrangement, extensive homoeologous gene loss, and widespread segmental allopolyploidy. Recent bursts of Copia mobility, not seen in other *Nicotiana* genomes, have probably aided *N. benthamiana*’s adaptation to a spectrum of Australian ecologies.

## Introduction

The genus *Nicotiana*, comprising ∼75 species, is predominantly endemic to the Americas and Australia^1^. Like most Solanaceae, it has a basic chromosome number of 12, with haploid DNA content ranging from 1.37Gb to 6.27Gb^2^. Section *Suaveolentes* (nicely smelling), includes *N. benthamiana* and is the largest allotetraploid group in the genus (∼35 species) with chromosome numbers ranging from 15 of 24, diagnostic of an allotetraplodization event followed by chromosome loss^3,4,5^ (Figure 1A). Almost all species in this section are indigenous to Australasia, which they apparently colonized during the Pliocene transition ∼5-6 MYA. The diploid ancestors of *N. benthamiana* most likely belonged to the *Sylvestres* and *Noctiflorae* sections, whose closest sequenced extant relatives are *N. sylvestris* (∼2.6Gb) and *N. glauca* (∼3.2Gb)^6–11^.

**Figure 1.**
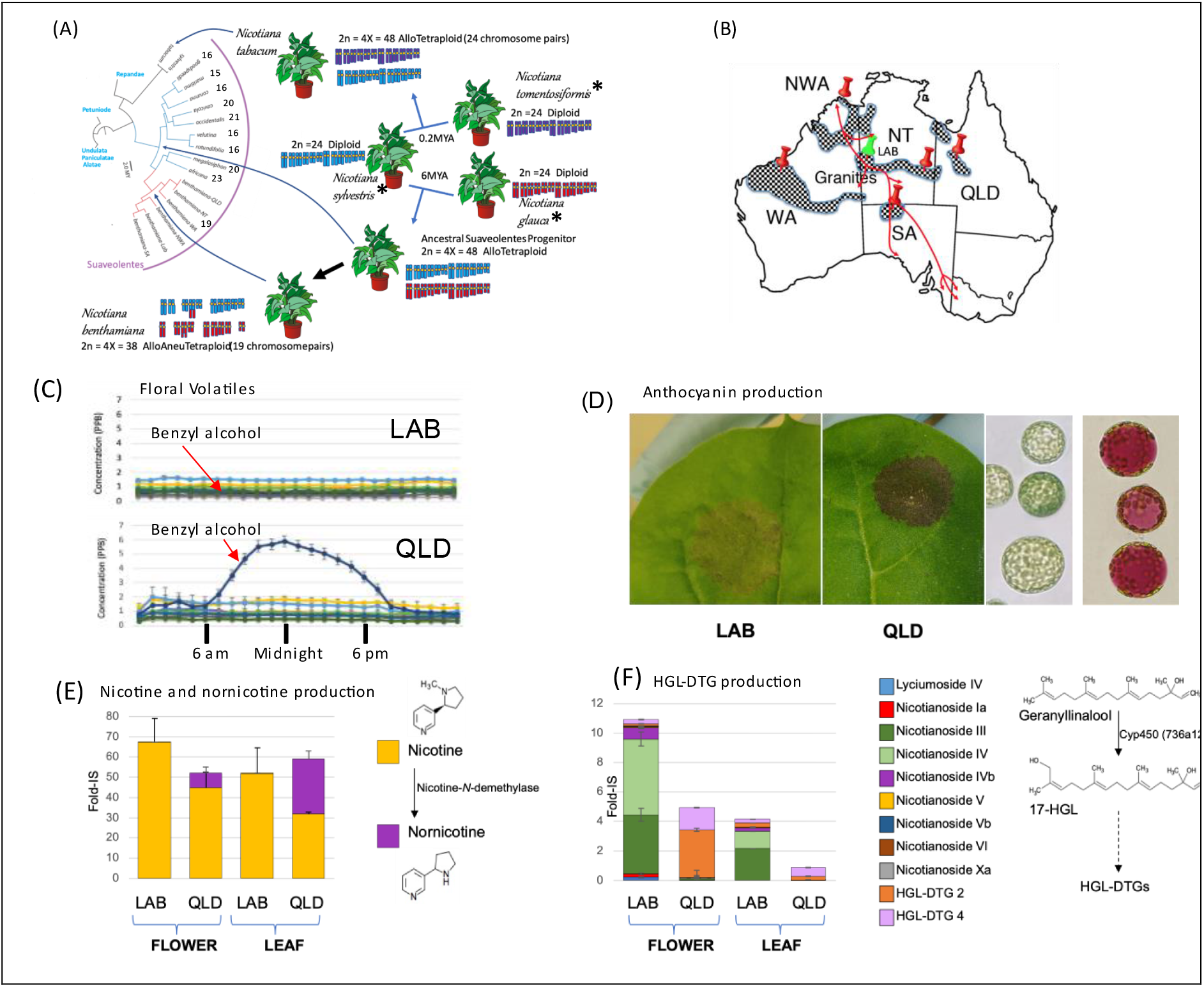
Phenotypic and biochemical diversity of *N. benthamiana*. **A**) Proposed phylogeny and origin of the *Suaveolentes* section compared to other Nicotianas. Chromosome numbers are indicated for each *Suaveolentes* species. Species highlighted by an asterisk are extant relatives of the putative parents of *N. benthamiana* and *N. tabacum*. B) Distribution of *N. benthamiana* in Australia (checkered regions). The physical locations of isolated *N. benthamiana* accessions reported in this study are shown by pins, and traditional indigenous trading routes are shown by red lines. C) Profiles of average emission of selected floral volatile compounds from LAB and QLD over a 24-hr period. Dark blue: Benzyl alcohol. For other compounds see Figure S1. D) Anthocyanin production 5 days after transient expression of AN-like MYB in LAB and QLD; right panels protoplasts isolated from control and AN-like infiltrated spots. E) Comparison of the accumulation of nicotine and nornicotine in flowers and leaves of LAB and QLD. The biochemical conversion of nicotine to nornicotine is shown on the right. F) Comparison of the accumulation of HGL-DTGs in flowers and leaves of LAB and QLD. The biochemical pathway is shown on the right.

*N. benthamiana* is the most important plant platform for biopharmaceutical protein and vaccine production^7,12^ and has been instrumental for fundamental discoveries in RNA interference, plant– pathogen interactions, metabolic pathway engineering, functional genomics, synthetic biology and gene editing^13^. All of this work has relied on plants derived from one accession line (that we term LAB), that appears to have originated from a single collection near the Granites gold mine in central Australia^7,14,15^ (Figure 1B). Several additional ecotypes have recently been described ^7,14–16^.

In this paper, we report chromosome level assemblies, epigenomic and metabolomic information for the LAB strain and the distantly related QLD ecotype, as well as their single nucleotide polymorphisms (SNPs) with two additional LAB and four additional wild accessions, providing a reference for the genus *Nicotiana*, and details about the genetic diversity of *N. benthamiana*. Furthermore, we study the genomic basis for their morphological, metabolic, and reproductive diversity, providing a comprehensive resource for fundamental research and biotechnology.

## Results and discussion

### An additional *N. benthamiana* ecotype resource

The QLD wild accession exhibits many morphological, developmental and metabolic differences to LAB^7,14–16^, such as outcrossing flowers^7,14–16^, moth-attracting volatile production at night, and the robust capacity to produce anthocyanins (Figures 1C-D, S1, S2, Table S1). The levels of a range of metabolites such as phenolic acids, flavonoids, amino acid derivatives, and metabolites involved in defense responses ^17–20^, such as nornicotine and hydroxygeranyl-linalool diterpene glycosides (HGL-DTG), exhibit marked differences between LAB and QLD (Figures 1E-F, S3, Table S2). LAB exhibited a higher number of underexpressed/non-functional biosynthetic pathways than QLD, except for phenolic acids and HGL-DTGs.

QLD is easily inter-crossable with LAB and is as amenable as LAB to genetic transformation, transient gene expression and genome editing (Figures S2-S5). For these characteristics, and their genetic distance (Figure 1A), both LAB and QLD were chosen for chromosome level genome sequence assemblies.

### Genome sequencing, assembly, annotation and genetic diversity

Long and short sequence reads of the LAB and QLD accessions were assembled into 19 chromosomes for each genome (Online methods, Figure S6). The chromosomes ranged in size from 128 to 182 Mb, with total genome sizes of ∼2.8Gb (LAB) and ∼2.9Gb (QLD), of which 99% and 96% respectively anchored to chromosomes (Table S3). This represents ∼94% of the expected genome size estimated from cytological staining^2^. The assemblies were annotated (Online methods, Figure S6, Table S3) to 45,797 and 49,636 gene models, respectively. Approximately 87% of the gene models in LAB and 75% in QLD are fully supported by RNA-sequencing and 97.7% of LAB EST sequences^21–23^ mapped to the LAB genome CDS. According to several quality scores, including the LTR Assembly Index^24^, the LAB and QLD assemblies were well above the standard requirements of the Earth Biogenome Project^25,26^ (Table S4), and showed far higher contiguity than any previously published *Nicotiana* genome assembly (Table 1), making them good candidates for a reference *Nicotiana* genome.

**Table 1.** Genome assembly metrics of LAB and QLD compared with reference genomes. Various genome assembly quality criteria (L50, N50, BUSCO score) are used to compare *N. benthamiana* with the other available genomes. Yellow shading highlights the assembled *N. benthamiana* genomes (this study), pink, those genomes with chr level assemblies and blue, draft assemblies. The values in brackets below *N. tabacum* and *N. attenuata* are those obtained from scaffold data alone.

| Species/<br>Ecotype | Scaffolds<br>>500nt | Chr | L50 | N50<br>(Mb) | Assembled<br>Genome Length<br>(Gb) | BUSCO% Complete<br>v10 |
| --- | --- | --- | --- | --- | --- | --- |
| *N.benthamiana* LAB | 19 | 19 | 10 | 145 | 2.75 | C:98.1%[S:46.0%,D:52.1%] |
| *N.benthamiana* QLD | 19 | 19 | 10 | 141 | 2.72 | C:98.0%[S:47.5%,D:50.5%] |
| *Arabidopsis thaliana* | 5 | 5 | 3 | 23 | 0.12 | C:99.2%[S:98.7%,D:0.5%] |
| Potato (dihaploid) | 12 | 12 | 6 | 59 | 0.74 | C:98.4%[S:96.6%,D:1.8%] |
| Tomato | 12 | 12 | 6 | 61 | 0.72 | C:97.8%[S:96.8%,D:1.0%] |
| Eggplant | 12 | 12 | 5 | 76 | 0.83 | C:84.2%[S:82.7%,D:1.5%] |
| Tobacco Chromosomes<br>(Scaffolds) | 24<br>(942,183) | 24 | 9<br>(3,998) | 84<br>(0.22) | 1.74<br>(4.01) | C:82.6%[S:61.2%,D:21.4%]<br>(C:96.8%[S:24.3%,D:72.5%]) |
| Capsicum | 12 | 12 | 6 | 221 | 2.56 | C:74.8%[S:73.7%,D:1.1%] |
| *N.attenuata* | 12<br>(37,194) | 12 | 498<br>(1,627) | 66<br>(0.45) | 0.73<br>(2.09) | C:48.5%[S:47.4%,D:1.1%]<br>(C:98.1%[S:95.9%,D:2.2%]) |
| *Petunia axilaris* | 17,630 | 12 | 17,630 | 1.24 | 1.20 | C:98.2%[S:95.6%,D:2.6%] |
| *N.benthamiana* LAB (USA v1.0.1) | 52,890 | 19 | 1,718 | 0.44 | 2.49 | C:98.2%[S:45.8%,D:52.4%] |
| *Petunia inflata* | 35,907 | 12 | 35,907 | 0.88 | 1.17 | C:97.9%[S:91.6%,D:6.3%] |
| *N.benthamiana* LAB (AU v0.5) | 77,255 | 19 | 1,903 | 0.39 | 2.49 | C:97.6%[S:47.5%,D:50.1%] |
| *N.sylvestris* | 125,957 | 12 | 7,255 | 0.08 | 2.01 | C:95.1%[S:93.3%,D:1.8%] |
| *N.tomemtoformis* | 90,682 | 12 | 5,663 | 0.15 | 1.62 | C:94.4%[S:92.6%,D:1.8%] |
| *N.obtusifolia* | 20,758 | 12 | 2,189 | 0.05 | 3.50 | C:94.3%[S:92.3%,D:2.0%] |
| *N.otophora* | 420,047 | 12 | 14,141 | 0.03 | 2.32 | C:76.0%[S:74.3%,D:1.7%] |

Gene mapping (Table S5A) revealed that 72%, 92% and 89% of the *N. benthamiana* genes are orthologous to those in tomato, *N. attenuata*, and tobacco, respectively. Similar numbers were obtained by protein cluster analysis (Figure S7, Table S5B). There were ∼1000 and ∼3000 genes specific to LAB and QLD, respectively. Based on BUSCO scores and comparison of the predicted protein lengths with their *Arabidopsis* best hits, the LAB and QLD annotations are better than most *Nicotiana* and *Solanaceae* annotations (Table S5C, Figure S8). A total of 369 and 383 potential miRNA families, and expression of 59 and 57, were detected in LAB and QLD, respectively.

The previously described NT, SA, WA and NWA wild accessions^14^ (Figure 1B), as well as the extensively used GFP-expressing transgenic line (16c) produced in David Baulcombe’s laboratory^23,27^ (EU-LAB) and (USA-LAB) were re-sequenced and mapped onto the LAB and QLD assemblies, Table S6). Single Nucleotide Polymorphisms (SNP) frequencies^28^ were very low amongst the three LAB accessions (<25 SNPs/Mb), confirming that our LAB assembly is an excellent resource for worldwide *N. benthamiana* laboratory isolates; SNPs between the four wild ecotypes mirrored the previously calculated evolutionary relationships^14^ (Table S6) and were similar in range to those of 20 *Capsicum annuum* accessions^29^. SA and LAB, originally collected from geographically well separated locations, have close genetic similarity (∼51 SNPs/Mb). One possible explanation is that Pitjuri (a chewing tobacco mixture often containing dried *N. benthamiana* aerial tissue) exchanged along ancient aboriginal traditional trading routes (Figure 1B) has transported seed between these locations over the last 60,000 years. The annotated genomes of LAB and QLD, containing tracks describing gene models, SNPs with other *N. benthamiana* isolates, gene expression across five tissues, location and expression of pre-miRNAs, and the epigenetic landscapes, are available on an interactive WebApollo browser^30^ (https://apollo.nbenth.com/).

### Identification of homeologous chromosomes, subgenomes and chromosome loss

The genomes of most diploid Solanaceous species consist of 12 chromosome pairs (2x=2n=24) encoding about 35,000 genes^31^. *N. tabacum*, an allotetraploid formed about 0.2-0.4 MYA^8,9^ has 24 chromosome pairs (2n=4x=48) encoding ∼70,000 genes^32,33^. In the estimated 5-6MY since the hybridization event basal to the Australian *Nicotiana* clade, *N. benthamiana* has lost 5 chromosome pairs to give a genome of 2n=4x=38^4,5^ (Figure 1A).

A mapping approach, similar to that used to identify the subgenomic memberships of the *N. tabacum* chromosomes^32–34^, was applied to *N. benthamiana* and *N. tabacum* using sequences from the genomes of *N. sylvestris, N. glauca* and *N. tomentosiformis* (Figure 2A). This recapitulated the previous tobacco results but, as previously predicted^8,9^, did not differentiate the *N. benthamiana* chromosomes into a *N. glauca*- and a *N. sylvestris*-related subgenome. Therefore, we took a different approach to identify the functional subgenomes. Syntenic sequences and blocks of orthologous genes were compared both within the highly syntenic LAB and QLD genomes, with the *N. sylvestris-* and *N. tomentosifomis-*derived chromosome assemblies of *N. tabacum*^*32*^ and with the *N. attenuata* genome assembly^34^ (Figure 2B). The chromosomal synteny between the LAB chromosomes and those of *N. sylvestris, N. tomentosiformis* and *N. attenuata* is apparent. A cladogram, derived from matrices of degrees of similarity of counterpart gene sequences of the *Nicotiana* set, clearly identified eight homeologous chromosome pairs and 3 orphan chromosomes (Figure 2C).

**Figure 2.**
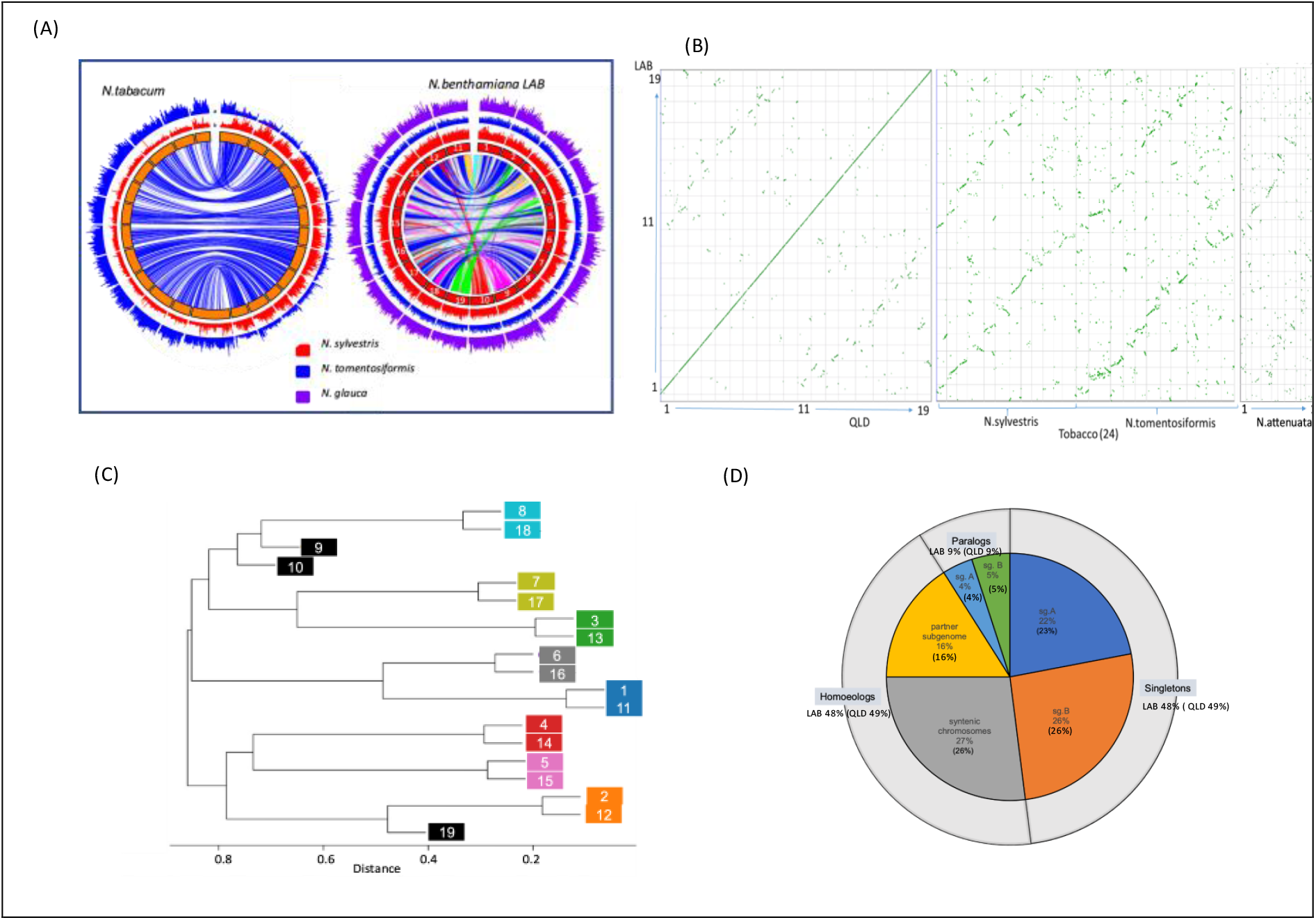
Subgenome and homeolog organization in *N. benthamiana*. **A)** The left hand Circos plot depicts the locations of the syntenic blocks (1Mbp) of *N. tomentosiformis* (blue) and *N. sylvestris* (red) on the *N. tabacum* genome, highlighting the subgenomes and their respective contribution to the subgenome structure of this species. The right hand Circos plot similarly locates the syntenic blocks of *N. tomentosiformis* (blue), *N. sylvestris* (red) and *N. glauca* (purple) on the *N. benthamiana* LAB genome, highlighting the difficulty in assigning ancestry for subgenomes in this species, which is characterised by extensive rearrangement of blocks between individual chromosomes. The lines in the centre join syntenic regions, highlighting the fragmentation of the *N. benthamiana* genome. **B)** Dot plot showing the relationship between the LAB and QLD chromosomes (central continuous line in the far-left panel) and the fragmented syntenic relationship between the subgenomes. Comparison of the *N. tabacum* genome consisting of 2 subgenomes with clear relationships to *N. sylvestris* and *N. tomentosiform*is revealed a fragmented relationship with *N. benthamiana* chromosomes. **C)** Cladogram highlighting the chromosome pairs and the 3 orphan chromosomes (9, 10 and 19). See text for further details. **D)** Retention and relocation of homeologous genes in *N. benthamiana* LAB and QLD genomes. Percentage values outside and within brackets are those for LAB and QLD, respectively, and show that about half of the original homeologous pairs have lost one member.

To separate the genome into two functional subgenomes we took a disjoint subset partitioning approach, enabled by the ∼50% of genes for which homeologous gene pairs were identified to be on chromosomes other than their predicted homeologous counterpart. Every combination of LAB chromosomes was assigned to two disjoint subsets and measured for the number of homeologous gene pairs distributed 1:1 between the two subsets. The best combination, excluding the genes on the three orphan chromosomes, gave a distribution of 8,543 gene pairs in opposite subgenomes and 1,999 gene pairs in the same subgenome (Tables S7A-B, Figure 2D). Visual comparison of *N. benthamiana* subgenomes with genomes of six other Solanaceous species using SynVisio^35^ revealed remarkable long range synteny across the family, especially in chromosomes 1, 2, 3 and 4, but still discernible in *N. tabacum* up to chr 7 (Figure 3A-B). By contrast, in *N. benthamiana* this synteny was abruptly interrupted after chromosome 4 (Figure 3B, 3E), probably due to the high degree of chromosomal rearrangements specific to this species. This endorsed the observed conservation of gene blocks in chromosomes 1 to 4 across all of the species.

**Figure 3.**
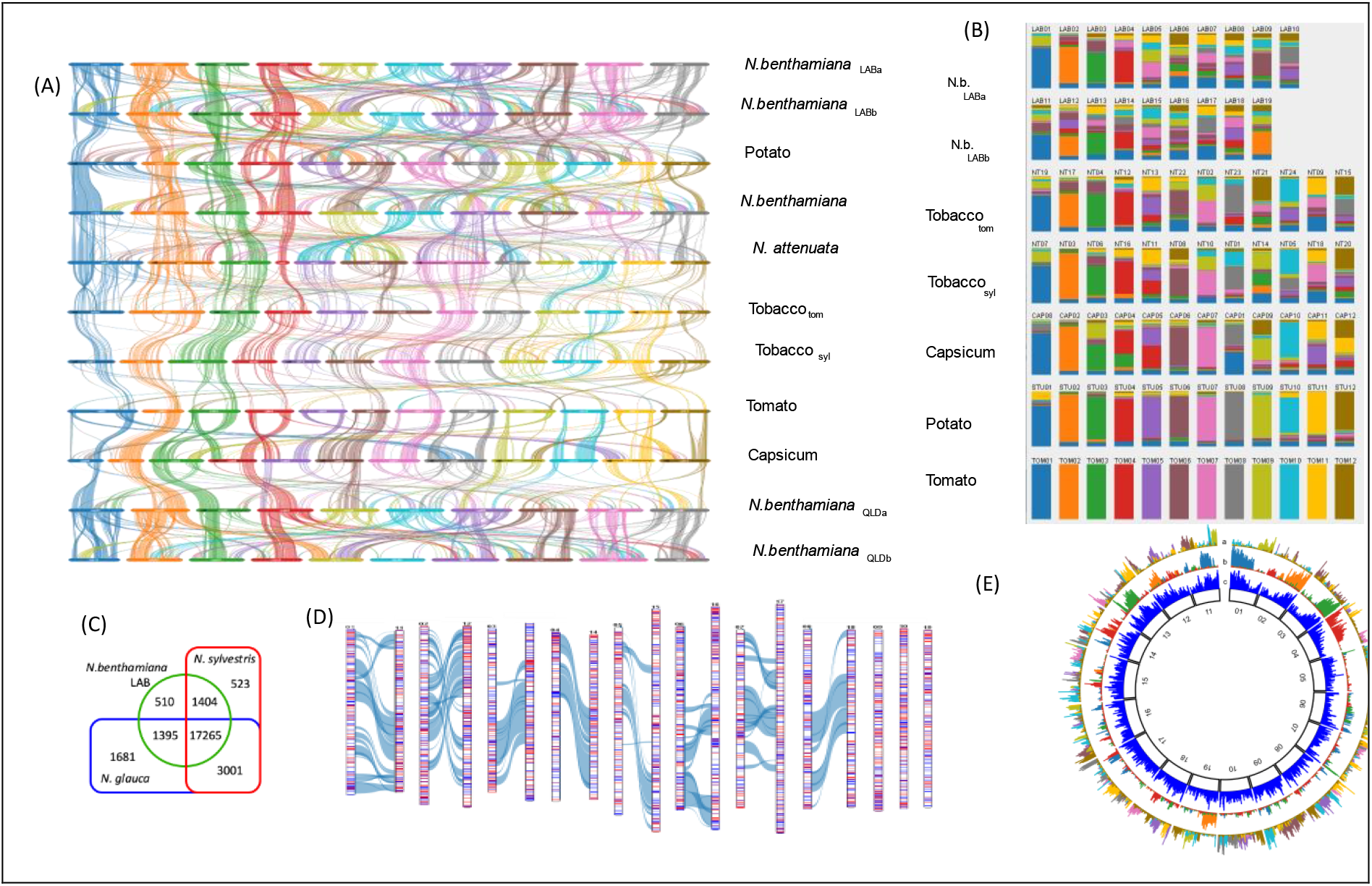
Gene block conservation across the Solanaceae and segmental allopolyploidisation in *N. benthamiana*. **A)** Waterfall plot showing the syntenic relationships between LAB, QLD and other related species as generated by SynVisio (https://synvisio.github.io/#/). **B)** Fraction of orthologous gene clusters in different Solanaceae chromosomes, highlighting a high conservation of chromosomes 1-4, and a declining conservation of remaining chromosomes; chromosome numbering follows the tomato-potato system. **C)** A Gibson Venn diagram showing the number of gene family clusters that are shared amongst LAB, *N. sylvestris* and *N. glauca*. **D)** Overlay of *N. glauca* (blue) and *N. sylvestris* (red) orthologous genes on LAB chromosomes; Grey/blue lines connecting chromosomes link syntenic blocks among the matched subgenome chromosomes. **E)** Circos plot of the physical distribution of syntenic blocks of tomato chromosomes 9-12 overlaid onto the LAB genome (track a), showing extensive fragmentation across the remaining LAB chromosomes. In contrast, an overlay of the syntenic blocks of Tomato chromosomes 1-4 onto the LAB genome clearly demonstrates a conservation of both sequence and location (track b). Track c shows the gene density across the LAB chromosomes.

The blocks of synteny between the two subgenomes of *N. benthamiana* are more numerous, larger and contiguous than with the *N. sylvestris-*derived subgenome of *N. tabacum* (Figure S9). To investigate this further, a cluster analysis was made using the proteomes predicted from our LAB assembly and from the available scaffold assemblies of *N. sylvestris* and *N. glauca* (Figure 3C). The LAB genes identified as clustering with *N. sylvestris* but not *N. glauca* genes, and vice-versa, were mapped onto the LAB genome (Figure 3D). This revealed that, even in the gene-rich, large, Solanaceae-wide syntenic blocks, extensive recombination has occurred between the two ancestral subgenomes. This suggests that the present *N. benthamiana* genome is the result of extensive “duplication/deletion” homeologous recombination^36^, or of repeated hybridization amongst the derivative populations from the original allotetraploid *Nicotiana* at the base of the Suaveolentes. These processes have produced chromosomes composed of genes from both ancestral parents, explaining the greater synteny between *N. benthamiana’s* homeologous chromosomes than with their *N. sylvestris* counterparts. This is also the likely cause of the low level of subgenome dominance (Figure S10). Subgenomes A and B encode 23,408 and 22,388 genes, respectively, and the overall transcript abundance of homeologs differs by only 1%, suggesting that the genome is in balanced but fluid harmony.

### Expansion and contraction of Transposable Elements

Polyploidization is often accompanied by bursts of transposable element (TE) activity^37–40^ and TEs, especially the type 1 long terminal repeat class (LTRs) such as Gypsy, are highly abundant in *Nicotiana*^*34*^. While Gypsy proliferation is obvious in the *N. benthamiana* genome, its content (∼1.5Gb) is more similar in size to those of the diploid *Nicotiana* species than to the allotetraploid *N. tabacum* or the combined sum of the extant ancestral parental diploid relatives, *N. glauca* and *N. sylvestris* (Figure 4A). A similar expansion of Gypsy content is evident in the recently reported pepper genome and is one of the main causes for its increased size^41^. However, as a percentage of genome size, all of these *Nicotianas*, including *N. benthamiana*, are about 50% Gypsy sequence, suggesting that the decreased Gypsy content in *N. benthamiana* is due to whole chromosome loss rather than TE-mediated genome purging ^42,43^.

**Figure 4.**
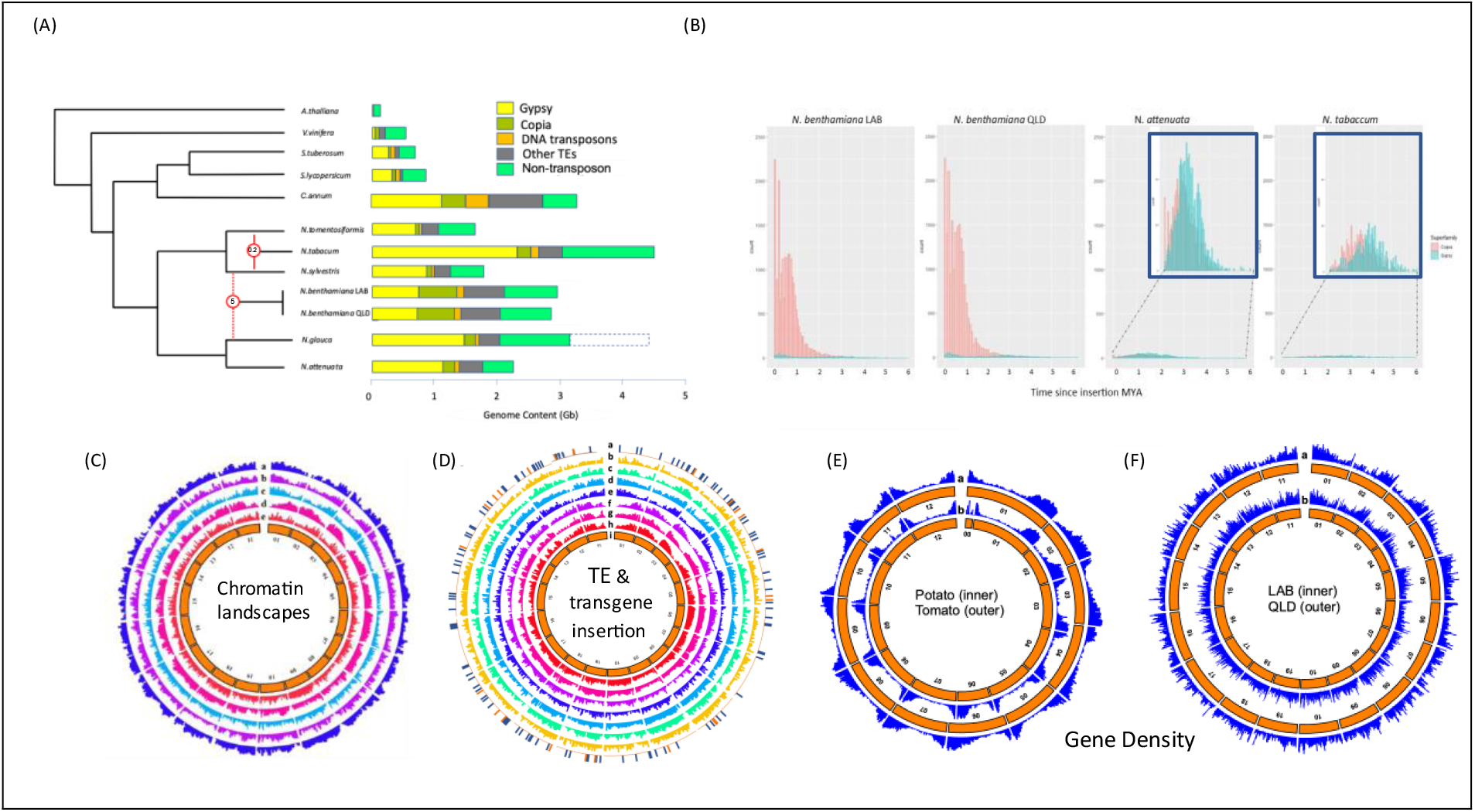
Transposon, epigenetic landscapes, and gene density of *N. benthamiana*. **A)** Relative complements of transposon and non-transposon content in *A. thaliana, V. vinifera* and key Solanaceous and *Nicotiana* species is presented as their calculated genome content in Gb. The dashed box for *N. glauca* indicates the genome size calculated from k-mer analysis (4.5Gb) while the composition of the genome is based on the current assembly of 3.2-3.5Gb. **B)** Estimated dates of LTR-retrotransposon insertion, calculated by sequence comparison between the LTRs of individual element insertions, in *N. benthamiana* LAB and QLD, compared to *N. attenuata* and *N. tabacum*. A clear and ongoing large burst of Copia element activity is evident in both LAB and QLD, that is absent in both *N. attenuata* and *N. tabacum*. The reported burst of Gypsy activity in *Nicotiana* species appears to predate the 6 MYA limit of our analysis. **C)** A Circos plot depicting the chromatin landscape as compared to gene content in LAB. Tracks a and b represent respectively the location of permissive histone marks H3K27ac and HsK4me3 within each LAB chr. Track c depicts the gene density across the LAB genome, while tracks d and e represent the location or repressive histone marks H3K9me2 and H3K27me3, respectively. As expected, there is a correlation between increasing gene density and permissive chromatin marks and reduction in gene density in gene poor chromosomal regions. **D)** Circos plot depicting the comparative locations of transgene insertions, LTR-retrotransposon insertion and methylation marks across LAB chromosomes. Track a; Transgene insertion sites; red ‘ticks’ represent insertions derived from stable transformation, blue ‘ticks’ represent insertions derived from transient agroinfiltration. Track b; Insertions of intact Copia TEs (i.e. containing matching LTRs and complete internal sequences). Track c; Insertion of all annotated Copia TEs, including fragmented elements. Track d; Distribution of CHH methylation marks. Track e; Gene density across the LAB genome. Track f; Insertions of all annotated Gypsy TEs, including fragmented elements. Track g; Distribution of CG methylation marks. Track h; distribution of CHG methylation marks. The innermost circle represents the numbered chromosomes. **E)** Distribution of gene densities on the chromosomes of potato and tomato. **F**) Distribution of gene densities on the chromosomes of LAB and QLD genomes.

Unlike any other sequenced Solanaceous species genome, including the closely related diploid *N. attenuata* and the polyploid *N. tabacum*, the *N. benthamiana* genome shows evidence of dramatic, recent Copia element proliferation (Figure 4A, B). Examining in more detail four different loci in the subgenomes of LAB and QLD and comparing them with their counterparts in tomato and other *Nicotianas* (Figures 5, S11, S12) revealed a common theme of expansion of intergenic regions in the *Nicotianas* compared to tomato, which, like in pepper, is largely due to Gypsy elements which are now highly fragmented. A second theme is tandem duplication in *Nicotiana*, followed by extensive pseudogenization specifically in *N. benthamiana*. An abundance of recent, intact Copia elements is also evident in *N. benthamiana*. Insertion dating (Figure 4B) reveals that sustained periods of Copia mobility started around 2 MYA, reaching a peak around 750 KY, and are still occurring. This coincides with the divergence of LAB and QLD, dated at ∼ 800 KYA^14^, and recently inserted Copia elements are evident in close proximity to key genes in all four loci that we examined (Figures 5, S11, S12) suggesting that the recent mobility has played a major role in the genome’s advancing diploidization and diversity. It is possible that the Copia explosion is common to all of the Australasian *Nicotianas* and, in conjunction with their allopolyploidy, this has fueled the adaptation enabling the widespread success of the *Suaveolentes* across some of the harshest climatic and ecological regions in Australia.

**Figure 5.**
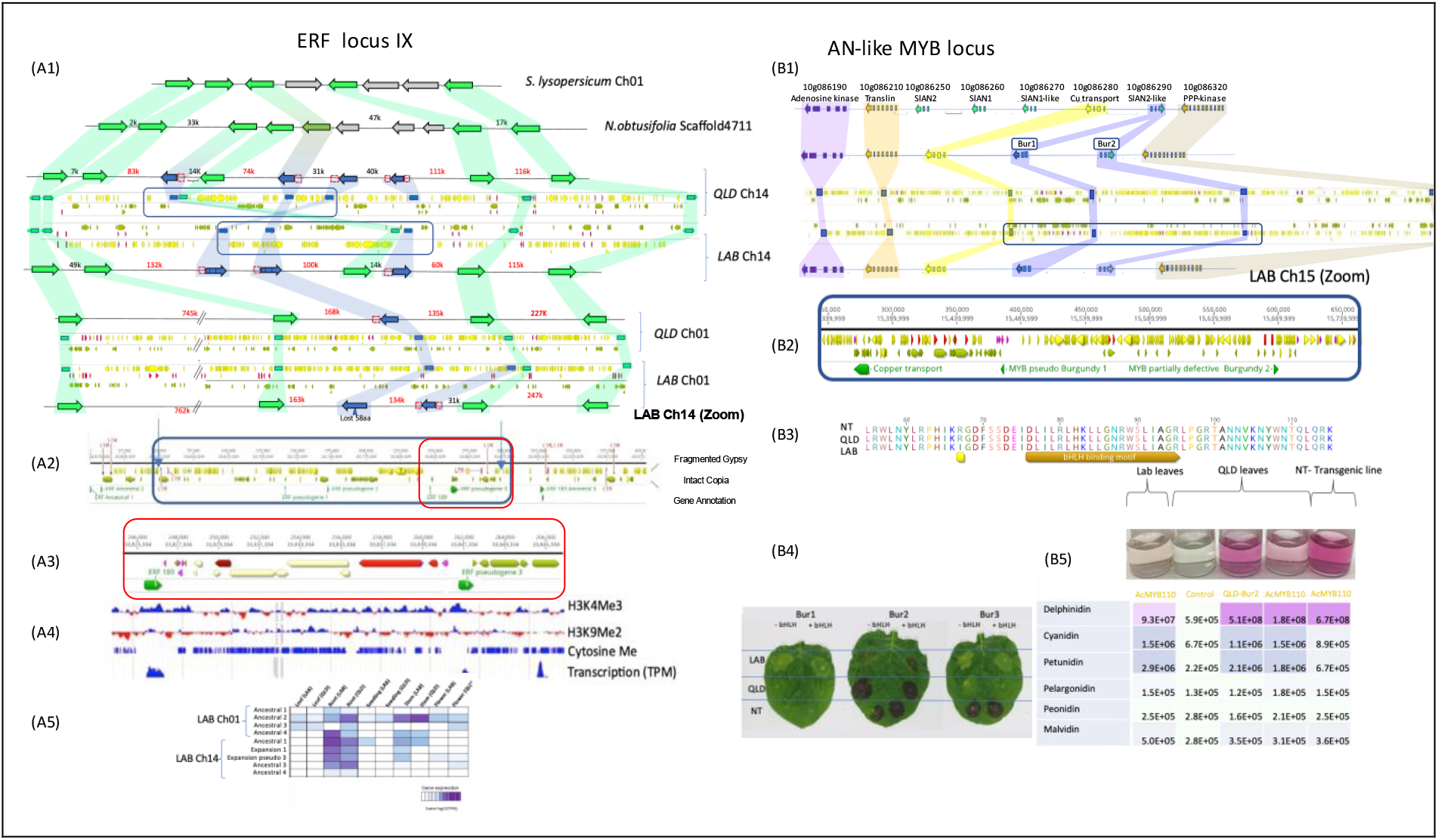
Comparison of ERF locus IX and AN-like MYB loci in LAB and QLD with other Solanaceae. **A1)** Synteny analysis of the ERF locus IX in tomato, *N. obtusifolia*, LAB and QLD shows lineage-specific tandem duplications of ERF189s, advanced diploidization through loss of gene function, and an inversion between LAB and QLD on chr 14, flanked by newly inserted Copia elements. Functional genes are shown in green; nonfunctional/pseudogenes are in blue. Gypsy, Copia and LTRs are indicated as yellow, olive green and red arrows respectively. Shading indicates the orthology relationships of ERF189 genes between different syntenic blocks. The inverted region of LAB chromosome 14 and the Gypsy and Copia landscape within the blue box is zoomed in the second panel (**A2**). The third panel (**A3**) is further zooming into the region indicated by a red box in (A2). The fourth panel (**A4**) depicts the epigenetic landscape (H3K4me3, H3K9me2 and cytosine methylation) and the expression of selected ERF189 genes in LAB. For H3K4me3, H3K9me2 enriched regions are shown in blue and the lack of the histone modification in red. Methylated cytosines are shown as blue bars. (**A5**) Tissue specific gene expression of Ancestral (the left-most and right-most two genes indicated in green in *N. obtusifolia*) and “ xpansion” (the three green genes in the middle of *N. obtusifolia*) genes. **B1)** Synteny analysis of the AN-like locus in tomato, LAB and QLD shows tandem duplication of SlAN2-like MYB genes in LAB and QLD with loss of gene function of 1 copy in QLD (Bur1) and both copies (Bur 1 & 2) in LAB. Loss of Bur2 in LAB is associated with a newly inserted Copia element. Functional genes are shown in bright green, nonfunctional/pseudogenes are in dark green. Gypsy, Copia and LTRs are indicated as yellow, olive green and red arrows respectively. Shading indicates the orthology relationships. The Gypsy and Copia landscape within the blue box are zoomed in the second panel (**B2**) The third panel (**B3**) shows the amino acid change in LAB Bur2 which alters its bHLH binding site. (**B4**) shows the function of Bur1 is defective in LAB, QLD and NT, that Bur2 is fully active in QLD and NT and may be partially restored by simultaneous over expression of bHLH in LAB. Bur3 is only functional in NT. (**B5**) Levels of different anthocyanins in LAB and QLD leaves following transient expression of AcMYB110 (an AN-like MYB from Kiwifruit) or QLD Bur2. For comparison the Anthocyanin levels were measured in NT stably transformed with an AcMYB110 construct.

### Epigenetic landscape and favoured sites for transgene integration

The epigenetic landscape of the LAB genome was examined for histone H3 methylation and acetylation, and cytosine methylation (Figures 4C-D, S13)^44^. Chromosomes 1,2,3,4,5, and to a lesser extent, 11 and 12, have a pronounced gradient of gene density across each chromosome which helps to reveal the correlation of high gene density with high levels of active histone marks (H3K4me3, H3K27ac). An inverse correlation of high gene density with repressive histone and DNA marks (H3K9me2 and CG and CHG methylation) is also apparent. These epigenetically repressed regions contain high levels of fragmented Gypsy elements whereas the active regions correlate with increased levels of intact Copia elements. The associations are also visible in the other chromosomes at a more localised level. The remarkably high level of recent Copia element insertions into regions with high gene density and active histone marks also correlates with high levels of CHH methylation which are likely driven by active transcription of these TEs.

To investigate whether epigenetic landscape has an influence on transgene insertion in the *N. benthamiana* genome, stable transgenic lines and leaf patches agro-infiltrated with transgene-encoding constructs were analysed for their insertion locations. From 40 independent transgenic lines, 23 sites could be mapped, and whole genome sequencing of the infiltrated patches identified 144 integration sites (Figure 4E). When adjusted for chromosome size, there was no significant bias for integration into any specific chromosome (p=0.19). However, integration into the gene body and promoter elements was more frequent than random and those inserting into intergenic regions were significantly closer to the gene borders. Transgene insertion into the gene body exhibited a much higher rate in transiently agroinfiltrated tissue than in stable transgenic lines, presumably because insertion-mediated dysfunctionality of some genes prevents whole plant development but is not lethal in tissue culture. The average intergenic size for *N. benthamiana* is ∼60Kb and the majority of transgenes inserted within the 10Kb region adjacent to a gene. A similar bias is apparent for active copies of both Copia and Gypsy (Figure 4D). Coupled with the histone and cytosine methylation status data, this supports the notion that trangenes and TEs are more able to integrate into the open chromatin of genes and adjacent regions than into the condensed core of intragenic zones.

### Advanced diploidization and pathway dysfunction in *N. benthamiana*

The loss of 5 chromosomes from the ancestral allotetraploid with retention of ∼50% of the genes in the genome as singletons (LAB sgA:10,075 sgB:11906; QLD sgA: 11,416 sgB:12,905) rather than homeologous pairs (Figure 2D, Tables S7A-B), indicates a loss of ∼20K genes/genome over 5MY. This complies with the estimation that the ancestral allotetraploid genome had ∼70K genes^31,32^ and, coupled with LAB’s genetic dysfunctions, explains the simple 3:1 Mendelian inheritance ratios of many traits in LAB x QLD crosses, such as virus susceptibility (*RDR1*)^14^, nor-nicotine production (*CYP82E*) and anthocyanin competence (*BUR2*). In each of these, LAB has the dysfunctional genes and pathways compared to QLD, and in the latter two loci of both ecotypes tandem gene duplication is evident in *Nicotianas*, but with progressive gene dysfunction in *N. benthamiana* (Figures 5A-B, S11, S12); a similar advancing diploidization is apparent in the ERF IX locus, encoding the transcription factors regulating nicotine production (Figure 4A), and in all these loci there is evidence of recently inserted Copia elements, suggestive of their role in the process. Diploid *Solanum* genomes and many non-solanaceous species exhibit high gene density bias towards the chromosome termini (Figure 4E-F). Interestingly, *N. benthamiana* chromosomes, especially 5-10 and 15-19, have a more uniform density. This unusual arrangement was likely caused by their formation through abundant inter-chromosomal recombination and by gene density dilution through the favoured insertion of TEs into the active chromatin of gene-rich regions.

### Conclusions and outlook

The exponential adoption of *Nicotiana benthamiana* as a model plant over the last two decades has produced vast amounts of data describing its responses to a wide spectrum of biotic and abiotic challenges, and this seems likely to continue unabated. Its use as a bioplatform to produce therapeutics has a similar trajectory. This dual role as model species and non-food bioproduction platform, on top of the unmatched capacity for fast transient transgene analysis, has made *N. benthamiana* the chassis of choice for testing and implementing the most advanced engineering approaches in Plant Synthetic Biology^45–47^. We produced a high-quality genome assembly of the LAB strain of *N. benthamiana* with fully annotated gene models, microRNA units, transposable elements, epigenetic landscapes, and chromosomal subgenomic membership, and made this publicly available on an interactive web-based genome browser. This enables decades of previously obtained data to be placed into a broader context, provides an important aid for future research and biotechnology, and facilitates involvement of the scientific community to expand and refine the resource.

Interestingly, *N. benthamiana* shows a recent explosion of Copia mobility and rapidly advancing diploidization. These two phenomena may or may not have a cause-effect relationship, but are apparently unique to this species, among sequenced *Nicotianas*, making it an excellent model species to study the course of diploidization and the dynamic balance of two subgenomes undergoing this process.

LAB is defective in many pathways compared to QLD, making it an excellent tool with which to explore interactions of transgene products, often from different species, in a host with reduced complexity. However, this also reduces the potential spectrum of metabolic and biotechnological engineering and possibly the comparability of the findings to fully endowed crop species. The high-quality genome assembly of QLD with its additional pathways and ∼ 3,000 genes, and the details about genomic diversity of an additional 4 wild and 2 laboratory isolates, provide resources to greatly enhance metabolic, developmental, and evolutionary studies. This is not only relevant to *N. benthamiana* but also across the Solanaceae, as it brings the genome of a *Nicotiana* species to the same chromosomal level of completeness (>95%) as tomato, eggplant, potato, and pepper.

## Acknowledgments

Work supported by the uropean ommission Horizon 2020 program, project “Developing Multipurpose Nicotiana Crops for Molecular Farming using New Plant Breeding Techniques (NEWCOTIANA)”, Grant Agreement n. 6 331.

## References

1. Knapp, S., Bohs, L., Nee, M. & Spooner, D. M. Solanaceae—A Model for Linking Genomics with Biodiversity. Comparative and Functional Genomics vol. 5 285–291 Preprint at https://doi.org/10.1002/cfg.393 (2004).

2. Narayan, R. K. J. Nuclear DNA changes, genome differentiation and evolution in Nicotiana (Solanaceae). Plant Syst. Evol. 157, 161–180 (1987).

3. Clarkson, J. J., Kelly, L. J., Leitch, A. R., Knapp, S. & Chase, M. W. Nuclear glutamine synthetase evolution in Nicotiana: phylogenetics and the origins of allotetraploid and homoploid (diploid) hybrids. Mol. Phylogenet. Evol. 55, 99–112 (2010).

4. Marks, C. E., Ladiges, P. Y. & Newbigin, E. Karyotypic variation in Nicotiana section Suaveolentes. Genet. Resour. Crop Evol. 58, 797–803 (2011).

5. Bally, J. et al. Nicotiana paulineana, a new Australian species in Nicotiana section Suaveolentes. Aust. Syst. Bot. 34, 477–484 (2021).

6. Byrne, M. et al. Birth of a biome: insights into the assembly and maintenance of the Australian arid zone biota. Mol. Ecol. 17, 4398–4417 (2008).

7. Bally, J. et al. The Rise and Rise of Nicotiana benthamiana: A Plant for All Reasons. Annu. Rev. Phytopathol. 56, 405–426 (2018).

8. Schiavinato, M., Marcet-Houben, M., Dohm, J. C., Gabaldón, T. & Himmelbauer, H. Parental origin of the allotetraploid tobacco Nicotiana benthamiana. Plant J. 102, 541–554 (2020).

9. Schiavinato, M., Bodrug-Schepers, A., Dohm, J. C. & Himmelbauer, H. Subgenome evolution in allotetraploid plants. Plant J. 106, 672–688 (2021).

10. Khafizova, G., Dobrynin, P., Polev, D. & Matveeva, T. Whole-genome sequencing of Nicotiana glauca. bioRxiv 211482 (2017) doi:10.1101/211482.

11. Usade, B. et al. The genome and metabolome of the tobacco tree, Nicotiana glauca: a potential renewable feedstock for the bioeconomy. bioRxiv 351429 (2018) doi:10.1101/351429.

12. LeBlanc, Z., Waterhouse, P. & Bally, J. Plant-Based Vaccines: The Way Ahead? Viruses 13, (2020).

13. Waterhouse, P. M. & Helliwell, C. A. Exploring plant genomes by RNA-induced gene silencing. Nat. Rev. Genet. 4, 29–38 (2003).

14. Bally, J. et al. The extremophile Nicotiana benthamiana has traded viral defence for early vigour. Nat Plants 1, 15165 (2015).

15. Drapal, M., Enfissi, E. M. A. & Fraser, P. D. Metabolic changes in leaves of N. tabacum and N. benthamiana during plant development. J. Plant Physiol. 265, 153486 (2021).

16. Drapal, M., Enfissi, E. M. A. & Fraser, P. D. Metabolic effects of agro-infiltration on N. benthamiana accessions. Transgenic Res. 30, 303–315 (2021).

17. Steppuhn, A., Gase, K., Krock, B., Halitschke, R. & Baldwin, I. T. Nicotine’s defensive function in nature. PLoS Biol. 2, E217 (2004).

18. de Boer, G. & Hanson, F. E. Feeding responses to solanaceous allelochemicals by larvae of the tobacco hornworm, Manduca sexta. Entomol. Exp. Appl. 45, 123–131 (1987).

19. Snook, M. E. et al. Hydroxygeranyllinalool Glycosides from Tobacco Exhibit Antibiosis Activity in the Tobacco Budworm [Heliothis virescens (F.)]. J. Agric. Food Chem. 45, 2299–2308 (1997).

20. Jassbi, A. R., Zamanizadehnajari, S., Kessler, D. & Baldwin, I. T. A New Acyclic Diterpene Glycoside from Nicotiana attenuata with a Mild Deterrent Effect on Feeding Manduca sexta Larvae. Zeitschrift für Naturforschung B 61, 1138–1142 (2006).

21. SGN EST search result - sol genomics network. https://solgenomics.net/search/est.pl?request_id=SGN-E1214852&request_from=0&request_type=automatic&search=Search.

22. Nakasugi, K., Crowhurst, R., Bally, J. & Waterhouse, P. Combining transcriptome assemblies from multiple de novo assemblers in the allo-tetraploid plant Nicotiana benthamiana. PLoS One 9, e91776 (2014).

23. Ruiz, M. T., Voinnet, O. & Baulcombe, D. C. Initiation and maintenance of virus-induced gene silencing. Plant Cell 10, 937–946 (1998).

24. Ou, S., Chen, J. & Jiang, N. Assessing genome assembly quality using the LTR Assembly Index (LAI). Nucleic Acids Res. 46, e126 (2018).

25. Howe, K. et al. Significantly improving the quality of genome assemblies through curation. Gigascience 10, (2021).

26. Lawniczak, M. K. N. et al. Standards recommendations for the Earth BioGenome Project. Proc. Natl. Acad. Sci. U. S. A. 119, (2022).

27. Philips, J. G. et al. The widely used Nicotiana benthamiana 16c line has an unusual T-DNA integration pattern including a transposon sequence. PLoS One 12, e0171311 (2017).

28. Lorenc, M. T. et al. Discovery of Single Nucleotide Polymorphisms in Complex Genomes Using SGSautoSNP. Biology 1, 370–382 (2012).

29. Qin, C. et al. Whole-genome sequencing of cultivated and wild peppers provides insights into Capsicum domestication and specialization. Proc. Natl. Acad. Sci. U. S. A. 111, 5135–5140 (2014).

30. Dunn, N. A. et al. Apollo: Democratizing genome annotation. PLoS Comput. Biol. 15, e1006790 (2019).

31. Barchi, L. et al. A chromosome-anchored eggplant genome sequence reveals key events in Solanaceae evolution. Sci. Rep. 9, 11769 (2019).

32. Edwards, K. D. et al. A reference genome for Nicotiana tabacum enables map-based cloning of homeologous loci implicated in nitrogen utilization efficiency. BMC Genomics 18, 448 (2017).

33. Brockmöller, T. et al. Nicotiana attenuata Data Hub (NaDH): an integrative platform for exploring genomic, transcriptomic and metabolomic data in wild tobacco. BMC Genomics 18, 79 (2017).

34. Xu, S. et al. Wild tobacco genomes reveal the evolution of nicotine biosynthesis. Proc. Natl. Acad. Sci. U. S. A. 114, 6133–6138 (2017).

35. Bandi, V. & Gutwin, C. Interactive Exploration of Genomic Conservation. Graphics Interface Preprint at https://doi.org/10.20380/GI2020.09 (2020).

36. Gaeta, R. T. & Chris Pires, J. Homoeologous recombination in allopolyploids: the polyploid ratchet. New Phytol. 186, 18–28 (2010).

37. Grandbastien, M.-A. et al. Stress activation and genomic impact of Tnt1 retrotransposons in Solanaceae. Cytogenet. Genome Res. 110, 229–241 (2005).

38. Kim, S. et al. Genome sequence of the hot pepper provides insights into the evolution of pungency in Capsicum species. Nat. Genet. 46, 270–278 (2014).

39. Kuang, H. et al. Identification of miniature inverted-repeat transposable elements (MITEs) and biogenesis of their siRNAs in the Solanaceae: new functional implications for MITEs. Genome Res. 19, 42–56 (2009).

40. Naito, K. et al. Unexpected consequences of a sudden and massive transposon amplification on rice gene expression. Nature 461, 1130–1134 (2009).

41. Liao, Y. et al. The 3D architecture of the pepper genome and its relationship to function and evolution. Nat. Commun. 13, 3479 (2022).

42. Lee, S.-I. & Kim, N.-S. Transposable elements, and genome size variations in plants. Genomics Inform. 12, 87–97 (2014).

43. Devos, K. M., Brown, J. K. M. & Bennetzen, J. L. Genome size reduction through illegitimate recombination counteracts genome expansion in Arabidopsis. Genome Res. 12, 1075–1079 (2002).

44. An, J. et al. J-Circos: an interactive Circos plotter. Bioinformatics 31, 1463–1465 (2015).

45. Mitiouchkina, T. et al. Plants with genetically encoded autoluminescence. Nat. Biotechnol. 38, 944–946 (2020).

46. Brophy, J. A. N. et al. Synthetic genetic circuits as a means of reprogramming plant roots. Science 377, 747–751 (2022).

47. Bernabé-Orts, J. M. et al. A memory switch for plant synthetic biology based on the phage ϕC31 integration system. Nucleic Acids Res. 48, 3379–3394 (2020).

