## Supplementary material for "A multi-omic *Nicotiana benthamiana* resource for fundamental research and biotechnology": online methods

### Plant lines

*N.benthamiana* LAB, NT, SA, WA, QLD, and NWA accessions have been described previously^1^. The EuLAB isolate is the extensively used GFP-expressing transgenic line (16c) produced in David Baulcombe’s laboratory, Sainsbury Institute, UK^2,3^ and USA-LAB has been described previously^4^**.** Plants were grown in a custom soil mix (UQ23 supplemented with Osmocote® slow release fertiliser) under controlled environmental conditions at a constant temperature of 25 oC with a 16 hour photoperiod and 8 hours dark period.

### RNA-Seq

Total RNA was isolated from four tissues (leaf, flower, stem, root) and seedlings (10 days) of LAB (6 weeks) and QLD (7 weeks) at the same developmental stage using TRIzol™ Reagent according to the manufacturer’s instructions. Libraries were constructed in triplicate for each tissue using NEBNext® ultra™ RNA Library Prep Kit for Illumina®, size selected (average 300nt), and sequenced on an Illumina HiSeq 2000/2500 system to produce 150 bp paired end reads.

### Extraction and analysis of secondary metabolites from plant tissues

Flower, leaf, stem, and roots were sampled as described for RNAseq and two biological replicates (individual plants) of the same samples of LAB and QLD were used for the metabolic analysis. Tissues were freeze-dried and homogeneously grounded in liquid nitrogen.

Semi-polar fraction was extracted from lyophilized ground tissue (3 mg for flower and root, and 5 mg for leaf and stem tissues) with 75% Methanol/0.1% v/v Formic acid, spiked with 0.25 µg/mL formononetin (Sigma-Aldrich) as internal standard. Metabolites were extracted at room temperature by continuous agitation for 30 min in MM 400 at 20 Hz. Samples were centrifuged at 20,000g for 20 min, 0.6 ml of the supernatant was transferred into filter (PTFE) vials for LC/MS analysis (0.2 µm pore size). Two independent extractions and analysis were performed for each biological replicate. LC conditions have been already described^5^. Five µL of filtered extract was injected in the LC/HESI/MS system, using a Q-exactive mass spectrometer (ThermoFisher Scientific). The ionization was performed using the heated electrospray ionization (HESI) source, with nitrogen used as sheath and auxiliary gas, and set to 35 and 10 units, respectively. The capillary temperature was 250°C, the spray voltage was set to 3.5 kV, the probe heater temperature was 330°C, and the S-lens RF level was set at 50. The acquisition was performed with the FTMS mass range of 110-1,600 m/z both in positive and in negative ion mode, with the following parameters: resolution 70,000, microscan 1, AGC target 1e6, and maximum injection time 100. Dd-MS2 parameters were as follows: resolution 17,500, intensity threshold 4.0e4, AGC target 2e4, maximum IT 50 ms, TopN 5, stepped NCE 15, 25, 40. All the chemicals and solvents used during the entire procedure were of LC/MS grade (Chromasolv, Merck Millipore).

Metabolic diversity was evaluated by comparing the MS spectra (positive ion mode) using SIEVE software (Thermofisher Scientific)^5^. The LC/MS spectra were processed by comparing tissues from each ecotype, only metabolites accumulating to levels of >2 fold change and pval<0.05 between the two ecotypes were selected. Metabolites were identified based on accurate masses in full MS together with MS2 spectra and/or authentic standard, using KEGG, Metfrag and PubChem mass databases(ST3). Relative levels of accumulation of investigated metabolites were measured and normalized relative to DW and to the internal standard, to correct for extraction and injection variability, as previously described^5^.

### Whole Genome Sequencing

High molecular weight genomic DNA from leaves or leaf nuclei of *N. benthamiana* LAB and QLD ecotypes were extracted as described previously^6^ and used for whole genome sequencing (lllumina, PacBio and Oxford Nanopore; see Figure S8). Short and long read sequencing was contracted to the Central Analytical Research Facility (CARF), Queensland University of Technology (QUT-CARF). The quality of the assemblies was determined using Merqury software (version 1.3)^7^. LAI scores were determined using the annotation obtained from the EDTA TE annotation pipeline^8^ and using the LAI sub-package of the LTR-retriever^9^ package according to Ou et al. 2018^10^ (see also <https://github.com/oushujun/EDTA/wiki/Calculate-LAI-from-EDTA-GFF3-files>).

### Genome assembly

The assembly pipeline is summarized in Figure S8. LAB and QLD contigs were assembled using CANU (version 1.81)^11^ and SparseAssembler k-mer 77 (version 20160205)/DBG2OLC (version 20160205)/Racon (version 1.3.2)^12–14^, respectively. Bionano optical mapping^15^ gave 44 and 37 super scaffolds for LAB and QLD, respectively, with contiguity statistic N50 values of 122 and 130Mbp. Juicer (version 1.6)^16^ and 3D-DNA (branch 201008)^17^ were used to generate Hi-C data and pre-assembly files. HiC libraries were produced as described by Dong et al.^18^, sequenced using the Illumina platform, and the aligned fragments from Juicer were further refined using Juicebox (version 2.12)^19^ and Citrus (<https://github.com/anjiyuan/Citrus>) to produce chromosome level assemblies. LR_Gapcloser^20^ (version 1.1) was used to close gaps with long reads to complete our genome assemblies. Afterwards, both assemblies were polished with Illumina reads using Pilon^21^ (version 1.23). Finally, Mercury^7^ (version 1.3) was used to categorize assembly quality based on EBG^22^. First, k-mer for DNA illumina sequence was generated by running the tool with “meryl k=21 count output xxx.meryl xxx.fastq.gz” and then generate k-mer completeness and QV value with “merqury.sh xxx.meryl <gene fasta> <prefix-output>”.

### Gene annotation

HISAT2 (version 2.1.0)^23^ generated BAM files using pooled RNAseq data (leaf, root, stem and seed) and Scallop (version 0.10.5)^24^ used to identify transcripts from the pooled RNAseq data. Transdecoder (https://github.com/TransDecoder/TransDecoder/) identified the coding and UTR regions. AUGUSTUS (version 3.2.3)^25^ was used to predict all possible transcripts based on the genome sequence. Combining the two gene annotations^26^, gave 267,000 and 255,000 genes for LAB and QLD, respectively. To filter out low confidence predicted genes, coding sequences of all the predicted genes were BLAST-searched^27^ against the NCBI NR (non-redundant) gene database and Solanaceae plants (tomato, potato, *N. attenuata*, *N. tabacum*) with the “identity” parameter gradually reduced until the BUSCO (version 4.0.5)^28^ score did not increase. Identity values of 86% (LAB) and 83% (QLD) corresponded to maximum BUSCO scores. To simplify the gene annotation, only one isoform (the longest) was retained where there appeared to be overlapping genes. Supplementing these high confidence genes with those lost in the analysis but identified by Scallop gave 45,796 and 49,636 genes for LAB and QLD, respectively. Gene mapping was undertaken by BLAST searching Tomato (<https://solgenomics.net/ftp/tomato_genome/assembly/build_4.00/>, v4.0), Nicotiana attenuata (<https://www.ncbi.nlm.nih.gov/assembly/GCF_001879085.1/>, including scaffolds) and Nicotiana tabacum (<https://solgenomics.net/ftp/genomes/Nicotiana_tabacum/edwards_et_al_2017/>) genomes with the sequences of gene coding regions from the LAB genome . Default BLAST settings were used.

### Protein Cluster Analysis

### Orthofinder version 2.5.4^29^ (using default settings) identified orthologous relationships among LAB, QLD, identified *N.tabacum*, *N.sylvestris*, *N.tomentosiformis*, *N.glauca*, *A.thaliana, V. vinifera S. lycopersicum and S. tuberosum.* See Supplementary Table S7C for details about versions and URLs.

### TE annotation

The EDTA pipeline (version 2.0.0)^8^ (<https://github.com/oushujun/EDTA>); last accessed 22/09/2022) was used to annotate the repeat element space for LAB, QLD, *N.attenuata* and *N.tabacum* with the following initiating command:

*>EDTA.pl --genome <genome fasta> --species others --step all -u --sensitive 0 --anno 1 --threads 48*.

The annotation of the *N. tabacum* genome only made use of the chromosome assembly available from the Sol Genomics network (<https://solgenomics.net/organism/Nicotiana_tabacum/genome>; file Nitab-v4.5_genome_Chr_Edwards2017.fasta.gz). The -u flag generates a file (*EDTA_raw/LTR/*.pass.list), containing estimations of LTR insertion times from LTR-retriever^9^ a component part of the EDTA pipeline. The estimation of insertion time is based on the numbers of polymorphisms calculated between the LTR sequences of intact LTE-TEs. Due to the lack of an accurate estimation of the neutral mutation rate in *N. benthamiana*, the default rate was set to that calculated for rice: 1.3e-8 substitutions per base pair per year^8^.

### MicroRNA annotation

The mature microRNA sequences from 79 plant species (Table S7E) were retrieved from miRbase (release 21; <https://www.mirbase.org/>) and used to identify miRs in *N. benthamiana* using Bowtie (version 2.0)^30^. To avoid missing IsomiRs, possible mature miRNA sequences with one mismatch were also identified using miRPlant (version 6)^31^. The expression levels of each miR and its precursor transcript was calculated from pooled data of libraries of small RNA and RNAseq reads (from this and previous studies^32,33^).

SNP calling

All Illumina genomic paired-end reads from each ecotype were aligned to the LAB and QLD assemblies using bowtie2^34^ (version 2.3.5). Duplicates reads were removed from each BAM file with Picard toolkit's (<https://broadinstitute.github.io/picard/>) (version 2.19) MarkDuplicates (*picard -Xmx25g MarkDuplicates ASSUME_SORT_ORDER=coordinate REMOVE_DUPLICATES=true*), and SAMtools^35^ (version 1.10) was used to keep unique (*samtools view -Sb -q 40*) and proper pair-end reads (*samtools view -@ 1 -hb -f 0x2 -F 2316*). Each read ID in the BAM file was modified by adding the ecotype's id using generate_subset_BAM.py from the SGSautoSNP^36^ pipeline (version 2.001). Next, BAM files for each cultivar were merged using SAMtools to produce BAM files for LAB and QLD. Finally, The SGSautoSNP.py script was used with default parameters.

### Chip-Seq

Cross-linking, chromatin isolation, nuclei lysis, chromatin shearing, and immunoprecipitation were carried out as described by Ranawaka *et al*.^6^. Antibodies against two active histone marks, anti-histone-H3-tri-methyl-K4 (Abcam ab8580) and anti-histone-H3-acetyl-K27 (Abcam ab4729), and two repressive histone marks, anti-histone-H3-tri-methyl-K27 (Abcam ab6002) and anti-histone-H3-di-methyl-K9 (Diagenode C15410060), were used in the immunoprecipitation step to generate the genome-wide histone modification landscapes of LAB and QLD. Libraries (two replicates per histone modification and control input) were prepared using NEBNext® Ultra™ II DNA Library Prep Kit for Illumina (E7645S) as per manufacturer’s specifications. ChIP-seq libraries of H3K9me2 were sequenced at the Central Analytical Research Facility (CARF), Queensland University of Technology (QUT-CARF), using Illumina NextSeq® 500 with output of 75 bp paired end reads (TG NextSeq® 500/550 High Output Kit v2, 75 cycle, TG-160-2005). Libraries of H3K4me3, H3K27me3, and H3K27ac were sequenced at Novogene International Private Limited (Singapore) on the Illumina HiSeq® 2000/ 2500 system to produce 150 bp paired end reads and analysed using the Galaxy platform (<https://usegalaxy.org.au>)^37^. Paired end reads were aligned against LAB and QLD genome assemblies using bowtie2 (version 2.4.2) with default settings^30^. Alignments with MAPQ < 40 were discarded prior to downstream analyses to ensure homoeolog specificity and accuracy. The deepTool, bamCompare^38^, was used to quantify and visualise histone marks across genes.

### Whole Genome Bisulfite Sequencing

Whole genome bisulfite sequencing samples were prepared with genomic DNA extracted from the same tissues used for ChIP-seq. Leaf genomic DNA from three replicates was extracted using a DNeasy Plant Mini Kit (QIAGEN, 69104). The bisulfite conversion of the DNA was carried out using the EZ DNA Methylation-Gold™ kit (ZYMO, D5005), and the bisulfite-treated DNA libraries constructed using the Illumina TruSeq DNA sample prep kit, following the manufacturer’s instructions. The library preparation and the subsequent next-generation sequencing were completed by Novogene HK Company Limited (Hong Kong Subsidiary). Paired end read (150 bp) sequencing of the bisulfite-treated DNA libraries was performed using an Illumina HiSeqX system.

### Methylation analysis

The high-quality reads from WGBS samples were aligned to LAB and QLD genome assemblies using the default settings of Bismark program (Version 0.19.0)^39^. PCR duplicates were removed with the deduplicate_bismark implemented in the Bismark program (version 0.19.0). Reads were mapped to the non-methylated chloroplast genome as a control to calculate the sodium bisulfite conversion rate of unmethylated cytosines which was >99.9% for all replicates (three replicates from each LAB and QLD). The cytosine methylation level was calculated using the bismark_methylation_extractor in Bismark (version 0.19.0). The methylation ratio of cytosine was calculated as the number of methylated cytosines divided by the number of reads covering that position.

### Calculation of relative expression levels of A and B subgenome homeologs

The MCScanX toolkit ^40^ was used to identify intraspecies syntenic blocks using protein sequences and chromosomal locations of genes (evalue 1e-10, max-target-seqs 6, masking 1, max-hsps 1). SynVisio^41^, an interactive multiscale synteny visualisation tool for McScanX, was used to visualise the gene-level collinearity. Genes in syntenic blocks were identified as homeologs, and the genes that could not find their homoeologous partners were identified as singletons. The average TPM expression of genes in each tissue type was calculated (average expression per tissue). Then, using the average expression of each gene per tissue, the global expression across all tissues was calculated. Global expression > 0.5 TPM was used for downstream analysis. Values of this combined analysis were used to determine the relative expression of homeologs. The analysis focused on homeologs which had a 1:1 correspondence across the two homoeologous subgenomes. The homoeologous pairs were defined as expressed when the sum of the A and B subgenome homeologs was > 0.5 TPM. This filtration included duplicate pairs where only a single homeolog was expressed. To standardise the relative expression of homeologs, the absolute TPM for each gene within the duplicate pair was normalised as follows. A and B represent the genes corresponding to the A and B homeologs in pairs.

Relative Expression of A=TPM(A)/(TPM(A)+TPM(B))

Relative Expression of B= TPM(B)/(TPM(A)+TPM(B))

The Kruskal-Wallis test was performed to statistically determine the homoeolog expression bias between subgenomes. Overrepresentation analysis was conducted using Fisher’s Exact Test. All the genes in *N. benthamiana* were blasted, mapped, and annotated using the Blast2Go suite^42^ and used as the background for the overrepresentation analysis. Highly suppressed genes in both subgenomes were assessed. Genes with a p value < 0.05 were considered significantly overrepresented.

### Identification and phylogenetic analysis of ERF189, NBS-LRR RPM1-like, Anthocyanin R2R3 Myb and Nicotine demethylase CYP82E genes

*ERF189*, *NBS-LRR RPM1*-like, Anthocyanin *R2R3 Myb* and *CYP82* genes in *N. benthamiana* were identified based on sequence homology using *N. attenuata* protein sequences (<http://nadh.ice.mpg.de/NaDH/others/data>) as query sequences for the Tblastn function on Apollo (https://apollo.nbenth.com/). *N.attenuata* CYP82 (NiAv7g20333) was identified by sequence similarity to tobacco CYP82E4, a demonstrated nicotine demethylase gene^43^. Phylogenetic trees were built using the identified sequences and their available counterparts in other *Nicotiana* sp. ( *N. attenuata*, *N. tabacum*, *N. sylvestris*, *N. tomentosiformis*) aligned using Muscle (version 3.8)^44^ . The best nucleotide substitution model was estimated based on jModeltest2 (version 2.1)^45^ and a tree constructed for each gene family using MrBayes (version 3.2.6)^46^ .

### Transgene insertion analysis

*Agrobacterium tumefaciens* (GV3101) transformed with a 35s-GFP-OCS construct (pBEN0317) was infiltrated into 4- weeks- old *N. benthamiana* leaves. After five days, agroinfiltrated leaves were collected. Total genomic DNA was extracted using the ISOLATE II Plant DNA Kit Bioline (BIO-52070) and pooled before library preparation using TruSeq® DNA Library Prep Kits (FC-121-2001). Sequencing was performed using the Illumina® Hiseq 2000 platform. Paired end reads were mapped to pBEN0317 binary vector using Burrows-Wheeler (BWA-MEM) (Version 0.7)^47^. To determine the T-DNA integration events, all split reads that partially overlapped the T-DNA region’s left and right borders were extracted and BLASTed (BLASTN) against the *N. benthamiana* genome. Reads with a percentage identity higher than 85% and an E value less than 1x10-5 were selected as high confidence transgene integration sites. A different approach was used to identify the broken reads. Reads were initially mapped to the *N.benthamiana* genome and mapped reads who’s mate is unmapped were extracted using Samtools view^35^. The filtered bam file was converted to fastq using bedtools Convert BAM to FastQ^48^. Reads were then BLASTed to the pBEN0317 vector. The reads which mapped to vectors with an E value of less than 1x10-5 and more than a 100 bp alignment were then BLASTed to the *N. benthamiana* genome. Reads with high identity (>95%) and ~50% coverage was identified as integrated T-DNA into the plant genome. For the stable transformation analysis, leaf tissues were collected from 5 weeks old *N. benthamiana* stable transgenic independent lines generated using pFN117 (Cas9) and pUQC-GFP-(218). Genomic DNA was extracted following the Cetyltrimethylammonium bromide (CTAB) method. Nested, insertion-specific primers for the right borders (RB1, RB2, and RB3) of pFN117 and pUQC-GFP-(218)-A were designed. Arbitrary degenerate primers (AD primers) and High-Throughput Thermal Asymmetric Interlaced Polymerase Chain Reaction (ht-TAIL-PCR) program were adapted from^49^. Purified PCR products were directly Sanger sequenced using RB3 primer, and the insertion sites were identified through a BLASTN search against the *N. benthamiana* genome. The number of stable and transient T-DNA insertion sites that intersect gene body, promoter, terminator, and transposable elements were determined using the bedtools Intersect tool (version 2.30.0)^48^ and the length to the closest gene from the insertion site calculated using RnaChipIntegrator (Version 1.1.0) (<https://github.com/fls-bioinformatics-core/RnaChipIntegrator>). The z-score test for two population proportions was used to determine the significant difference between 10kb, 10-20kb, 20-30kb and 30-40kb intervals from all stable, transient transgene insertion sites and randomly selected sites in the *N. benthamiana* genome.

References

1.	Bally, J. *et al.* The extremophile Nicotiana benthamiana has traded viral defence for early vigour. *Nat Plants* **1**, 15165 (2015).

2.	Ruiz, M. T., Voinnet, O. & Baulcombe, D. C. Initiation and maintenance of virus-induced gene silencing. *Plant Cell* **10**, 937–946 (1998).

3.	Philips, J. G. *et al.* The widely used Nicotiana benthamiana 16c line has an unusual T-DNA integration pattern including a transposon sequence. *PLoS One* **12**, e0171311 (2017).

4.	Bombarely, A. *et al.* A draft genome sequence of Nicotiana benthamiana to enhance molecular plant-microbe biology research. *Mol. Plant. Microbe. Interact.* **25**, 1523–1530 (2012).

5.	Sulli, M. *et al.* An Eggplant Recombinant Inbred Population Allows the Discovery of Metabolic QTLs Controlling Fruit Nutritional Quality. *Front. Plant Sci.* **12**, 638195 (2021).

6.	Ranawaka, B., Tanurdzic, M., Waterhouse, P. & Naim, F. An optimised chromatin immunoprecipitation (ChIP) method for starchy leaves of Nicotiana benthamiana to study histone modifications of an allotetraploid plant. *Mol. Biol. Rep.* **47**, 9499–9509 (2020).

7.	Rhie, A., Walenz, B. P., Koren, S. & Phillippy, A. M. Merqury: reference-free quality, completeness, and phasing assessment for genome assemblies. *Genome Biol.* **21**, 245 (2020).

8.	Ou, S. *et al.* Benchmarking transposable element annotation methods for creation of a streamlined, comprehensive pipeline. *Genome Biol.* **20**, 275 (2019).

9.	Ou, S. & Jiang, N. LTR_retriever: A Highly Accurate and Sensitive Program for Identification of Long Terminal Repeat Retrotransposons. *Plant Physiol.* **176**, 1410–1422 (2018).

10.	Ou, S., Chen, J. & Jiang, N. Assessing genome assembly quality using the LTR Assembly Index (LAI). *Nucleic Acids Res.* **46**, e126 (2018).

11.	Koren, S. *et al.* Canu: scalable and accurate long-read assembly via adaptive*k*-mer weighting and repeat separation. Preprint at https://doi.org/[10.1101/071282](http://dx.doi.org/10.1101/071282).

12.	[Ye, C., Ma, Z. S., Cannon, C. H., Pop, M. & Yu, D. W. SparseAssembler: de novo Assembly with the Sparse de Bruijn Graph. *arXiv [cs.DS]* (2011).](http://paperpile.com/b/vUxBk4/epcta)

13.	Ye, C., Hill, C. M., Wu, S., Ruan, J. & Ma, Z. DBG2OLC: efficient assembly of large genomes using long erroneous reads of the third generation sequencing technologies. Sci Rep 6: 31900. *North Pacific individual showing demographic*.

14.	Vaser, R., Sović, I., Nagarajan, N. & Šikić, M. Fast and accurate de novo genome assembly from long uncorrected reads. *Genome Res.* **27**, 737–746 (2017).

15.	Liu, J. *et al.* Gapless assembly of maize chromosomes using long-read technologies. *Genome Biol.* **21**, 121 (2020).

16.	Durand, N. C. *et al.* Juicer Provides a One-Click System for Analyzing Loop-Resolution Hi-C Experiments. *Cell Syst* **3**, 95–98 (2016).

17.	Dudchenko, O. *et al.* De novo assembly of the genome using Hi-C yields chromosome-length scaffolds. *Science* **356**, 92–95 (2017).

18.	Dong, P. *et al.* 3D Chromatin Architecture of Large Plant Genomes Determined by Local A/B Compartments. *Mol. Plant* **10**, 1497–1509 (2017).

19.	Durand, N. C. *et al.* Juicebox Provides a Visualization System for Hi-C Contact Maps with Unlimited Zoom. *Cell Syst* **3**, 99–101 (2016).

20.	Xu, G.-C. *et al.* LR_Gapcloser: a tiling path-based gap closer that uses long reads to complete genome assembly. *Gigascience* **8**, (2019).

21.	Walker, B. J. *et al.* Pilon: an integrated tool for comprehensive microbial variant detection and genome assembly improvement. *PLoS One* **9**, e112963 (2014).

22.	Howe, K. *et al.* Significantly improving the quality of genome assemblies through curation. *Gigascience* **10**, (2021).

23.	Kim, D., Langmead, B. & Salzberg, S. L. HISAT: a fast spliced aligner with low memory requirements. *Nat. Methods* **12**, 357–360 (2015).

24.	Shao, M. & Kingsford, C. Accurate assembly of transcripts through phase-preserving graph decomposition. *Nat. Biotechnol.* **35**, 1167–1169 (2017).

25.	Stanke, M. & Morgenstern, B. AUGUSTUS: a web server for gene prediction in eukaryotes that allows user-defined constraints. *Nucleic Acids Research* vol. 33 W465–W467 Preprint at https://doi.org/[10.1093/nar/gki458](http://dx.doi.org/10.1093/nar/gki458) (2005).

26.	Dainat, J. AGAT: Another Gff Analysis Toolkit to handle annotations in any GTF/GFF format. *Version v0* **4**, 10–5281 (2020).

27.	Altschul, S. F., Gish, W., Miller, W., Myers, E. W. & Lipman, D. J. Basic local alignment search tool. *J. Mol. Biol.* **215**, 403–410 (1990).

28.	Manni, M., Berkeley, M. R., Seppey, M. & Zdobnov, E. M. BUSCO: Assessing Genomic Data Quality and Beyond. *Curr Protoc* **1**, e323 (2021).

29.	Emms, D. M. & Kelly, S. OrthoFinder: phylogenetic orthology inference for comparative genomics. *Genome Biol.* **20**, 238 (2019).

30.	Langmead, B. & Salzberg, S. L. Fast gapped-read alignment with Bowtie 2. *Nat. Methods* **9**, 357–359 (2012).

31.	An, J., Lai, J., Sajjanhar, A., Lehman, M. L. & Nelson, C. C. miRPlant: an integrated tool for identification of plant miRNA from RNA sequencing data. *BMC Bioinformatics* **15**, 275 (2014).

32.	Huen, A., Bally, J. & Smith, P. Identification and characterisation of microRNAs and their target genes in phosphate-starved Nicotiana benthamiana by small RNA deep sequencing and 5’RACE analysis. *BMC Genomics* **19**, 940 (2018).

33.	Baksa, I. *et al.* Identification of Nicotiana benthamiana microRNAs and their targets using high throughput sequencing and degradome analysis. *BMC Genomics* **16**, 1025 (2015).

34.	Langmead, B., Wilks, C., Antonescu, V. & Charles, R. Scaling read aligners to hundreds of threads on general-purpose processors. *Bioinformatics* **35**, 421–432 (2019).

35.	Li, H. *et al.* The Sequence Alignment/Map format and SAMtools. *Bioinformatics* **25**, 2078–2079 (2009).

36.	Lorenc, M. T. *et al.* Discovery of Single Nucleotide Polymorphisms in Complex Genomes Using SGSautoSNP. *Biology* **1**, 370–382 (2012).

37.	Afgan, E. *et al.* The Galaxy platform for accessible, reproducible and collaborative biomedical analyses: 2018 update. *Nucleic Acids Res.* **46**, W537–W544 (2018).

38.	Ramírez, F. *et al.* deepTools2: a next generation web server for deep-sequencing data analysis. *Nucleic Acids Res.* **44**, W160–5 (2016).

39.	Krueger, F. & Andrews, S. R. Bismark: a flexible aligner and methylation caller for Bisulfite-Seq applications. *Bioinformatics* **27**, 1571–1572 (2011).

40.	Wang, Y. *et al.* MCScanX: a toolkit for detection and evolutionary analysis of gene synteny and collinearity. *Nucleic Acids Res.* **40**, e49 (2012).

41.	Bandi, V. & Gutwin, C. Interactive Exploration of Genomic Conservation. (2022).

42.	Götz, S. *et al.* High-throughput functional annotation and data mining with the Blast2GO suite. *Nucleic Acids Res.* **36**, 3420–3435 (2008).

43.	Siminszky, B., Gavilano, L., Bowen, S. W. & Dewey, R. E. Conversion of nicotine to nornicotine in Nicotiana tabacum is mediated by CYP82E4, a cytochrome P450 monooxygenase. *Proc. Natl. Acad. Sci. U. S. A.* **102**, 14919–14924 (2005).

44.	Edgar, S. M. & Theriot, E. C. Phylogeny of Aulacoseira (Bacillariophyta) based on molecules and morphology1. *J. Phycol.* **40**, 772–788 (2004).

45.	Darriba, D., Taboada, G. L., Doallo, R. & Posada, D. jModelTest 2: more models, new heuristics and parallel computing. *Nat. Methods* **9**, 772 (2012).

46.	Ronquist, F. *et al.* MrBayes 3.2: efficient Bayesian phylogenetic inference and model choice across a large model space. *Syst. Biol.* **61**, 539–542 (2012).

47.	Li, H. & Durbin, R. Fast and accurate long-read alignment with Burrows-Wheeler transform. *Bioinformatics* **26**, 589–595 (2010).

48.	Quinlan, A. R. & Hall, I. M. BEDTools: a flexible suite of utilities for comparing genomic features. *Bioinformatics* **26**, 841–842 (2010).

49.	Singer, T. & Burke, E. High-throughput TAIL-PCR as a tool to identify DNA flanking insertions. *Methods Mol. Biol.* **236**, 241–272 (2003).

1.	Bally, J. et al. The extremophile Nicotiana benthamiana has traded viral defence for early vigour. Nat Plants 1, 15165 (2015).

2.	Ruiz, M. T., Voinnet, O. & Baulcombe, D. C. Initiation and maintenance of virus-induced gene silencing. Plant Cell 10, 937–946 (1998).

3.	Philips, J. G. et al. The widely used Nicotiana benthamiana 16c line has an unusual T-DNA integration pattern including a transposon sequence. PLoS One 12, e0171311 (2017).

4.	Bombarely, A. et al. A draft genome sequence of Nicotiana benthamiana to enhance molecular plant-microbe biology research. Mol. Plant. Microbe. Interact. 25, 1523–1530 (2012).

5.	Sulli, M. et al. An Eggplant Recombinant Inbred Population Allows the Discovery of Metabolic QTLs Controlling Fruit Nutritional Quality. Front. Plant Sci. 12, 638195 (2021).

6.	Ranawaka, B., Tanurdzic, M., Waterhouse, P. & Naim, F. An optimised chromatin immunoprecipitation (ChIP) method for starchy leaves of Nicotiana benthamiana to study histone modifications of an allotetraploid plant. Mol. Biol. Rep. 47, 9499–9509 (2020).

7.	Rhie, A., Walenz, B. P., Koren, S. & Phillippy, A. M. Merqury: reference-free quality, completeness, and phasing assessment for genome assemblies. Genome Biol. 21, 245 (2020).

8.	Ou, S. et al. Benchmarking transposable element annotation methods for creation of a streamlined, comprehensive pipeline. Genome Biol. 20, 275 (2019).

9.	Ou, S. & Jiang, N. LTR_retriever: A Highly Accurate and Sensitive Program for Identification of Long Terminal Repeat Retrotransposons. Plant Physiol. 176, 1410–1422 (2018).

10.	Ou, S., Chen, J. & Jiang, N. Assessing genome assembly quality using the LTR Assembly Index (LAI). Nucleic Acids Res. 46, e126 (2018).

11.	Koren, S. et al. Canu: scalable and accurate long-read assembly via adaptivek-mer weighting and repeat separation. Preprint at https://doi.org/10.1101/071282.

12.	Ye, C., Ma, Z. S., Cannon, C. H., Pop, M. & Yu, D. W. SparseAssembler: de novo Assembly with the Sparse de Bruijn Graph. arXiv [cs.DS] (2011).

13.	Ye, C., Hill, C. M., Wu, S., Ruan, J. & Ma, Z. DBG2OLC: efficient assembly of large genomes using long erroneous reads of the third generation sequencing technologies. Sci Rep 6: 31900. North Pacific individual showing demographic.

14.	Vaser, R., Sović, I., Nagarajan, N. & Šikić, M. Fast and accurate de novo genome assembly from long uncorrected reads. Genome Res. 27, 737–746 (2017).

15.	Liu, J. et al. Gapless assembly of maize chromosomes using long-read technologies. Genome Biol. 21, 121 (2020).

16.	Durand, N. C. et al. Juicer Provides a One-Click System for Analyzing Loop-Resolution Hi-C Experiments. Cell Syst 3, 95–98 (2016).

17.	Dudchenko, O. et al. De novo assembly of the genome using Hi-C yields chromosome-length scaffolds. Science 356, 92–95 (2017).

18.	Dong, P. et al. 3D Chromatin Architecture of Large Plant Genomes Determined by Local A/B Compartments. Mol. Plant 10, 1497–1509 (2017).

19.	Durand, N. C. et al. Juicebox Provides a Visualization System for Hi-C Contact Maps with Unlimited Zoom. Cell Syst 3, 99–101 (2016).

20.	Xu, G.-C. et al. LR_Gapcloser: a tiling path-based gap closer that uses long reads to complete genome assembly. Gigascience 8, (2019).

21.	Walker, B. J. et al. Pilon: an integrated tool for comprehensive microbial variant detection and genome assembly improvement. PLoS One 9, e112963 (2014).

22.	Howe, K. et al. Significantly improving the quality of genome assemblies through curation. Gigascience 10, (2021).

23.	Kim, D., Langmead, B. & Salzberg, S. L. HISAT: a fast spliced aligner with low memory requirements. Nat. Methods 12, 357–360 (2015).

24.	Shao, M. & Kingsford, C. Accurate assembly of transcripts through phase-preserving graph decomposition. Nat. Biotechnol. 35, 1167–1169 (2017).

25.	Stanke, M. & Morgenstern, B. AUGUSTUS: a web server for gene prediction in eukaryotes that allows user-defined constraints. Nucleic Acids Research vol. 33 W465–W467 Preprint at https://doi.org/10.1093/nar/gki458 (2005).

26.	Dainat, J. AGAT: Another Gff Analysis Toolkit to handle annotations in any GTF/GFF format. Version v0 4, 10–5281 (2020).

27.	Altschul, S. F., Gish, W., Miller, W., Myers, E. W. & Lipman, D. J. Basic local alignment search tool. J. Mol. Biol. 215, 403–410 (1990).

28.	Manni, M., Berkeley, M. R., Seppey, M. & Zdobnov, E. M. BUSCO: Assessing Genomic Data Quality and Beyond. Curr Protoc 1, e323 (2021).

29.	Emms, D. M. & Kelly, S. OrthoFinder: phylogenetic orthology inference for comparative genomics. Genome Biol. 20, 238 (2019).

30.	Langmead, B. & Salzberg, S. L. Fast gapped-read alignment with Bowtie 2. Nat. Methods 9, 357–359 (2012).

31.	An, J., Lai, J., Sajjanhar, A., Lehman, M. L. & Nelson, C. C. miRPlant: an integrated tool for identification of plant miRNA from RNA sequencing data. BMC Bioinformatics 15, 275 (2014).

32.	Huen, A., Bally, J. & Smith, P. Identification and characterisation of microRNAs and their target genes in phosphate-starved Nicotiana benthamiana by small RNA deep sequencing and 5’RACE analysis. BMC Genomics 19, 940 (2018).

33.	Baksa, I. et al. Identification of Nicotiana benthamiana microRNAs and their targets using high throughput sequencing and degradome analysis. BMC Genomics 16, 1025 (2015).

34.	Langmead, B., Wilks, C., Antonescu, V. & Charles, R. Scaling read aligners to hundreds of threads on general-purpose processors. Bioinformatics 35, 421–432 (2019).

35.	Li, H. et al. The Sequence Alignment/Map format and SAMtools. Bioinformatics 25, 2078–2079 (2009).

36.	Lorenc, M. T. et al. Discovery of Single Nucleotide Polymorphisms in Complex Genomes Using SGSautoSNP. Biology 1, 370–382 (2012).

37.	Afgan, E. et al. The Galaxy platform for accessible, reproducible and collaborative biomedical analyses: 2018 update. Nucleic Acids Res. 46, W537–W544 (2018).

38.	Ramírez, F. et al. deepTools2: a next generation web server for deep-sequencing data analysis. Nucleic Acids Res. 44, W160–5 (2016).

39.	Krueger, F. & Andrews, S. R. Bismark: a flexible aligner and methylation caller for Bisulfite-Seq applications. Bioinformatics 27, 1571–1572 (2011).

40.	Wang, Y. et al. MCScanX: a toolkit for detection and evolutionary analysis of gene synteny and collinearity. Nucleic Acids Res. 40, e49 (2012).

41.	Bandi, V. & Gutwin, C. Interactive Exploration of Genomic Conservation. (2022).

42.	Siminszky, B., Gavilano, L., Bowen, S. W. & Dewey, R. E. Conversion of nicotine to nornicotine in Nicotiana tabacum is mediated by CYP82E4, a cytochrome P450 monooxygenase. Proc. Natl. Acad. Sci. U. S. A. 102, 14919–14924 (2005).

43.	Li, H. & Durbin, R. Fast and accurate long-read alignment with Burrows-Wheeler transform. Bioinformatics 26, 589–595 (2010).

44.	Quinlan, A. R. & Hall, I. M. BEDTools: a flexible suite of utilities for comparing genomic features. Bioinformatics 26, 841–842 (2010).

45.	Singer, T. & Burke, E. High-throughput TAIL-PCR as a tool to identify DNA flanking insertions. Methods Mol. Biol. 236, 241–272 (2003).
