## supplementary figures for "A multi-omic *Nicotiana benthamiana* resource for fundamental research and biotechnology"

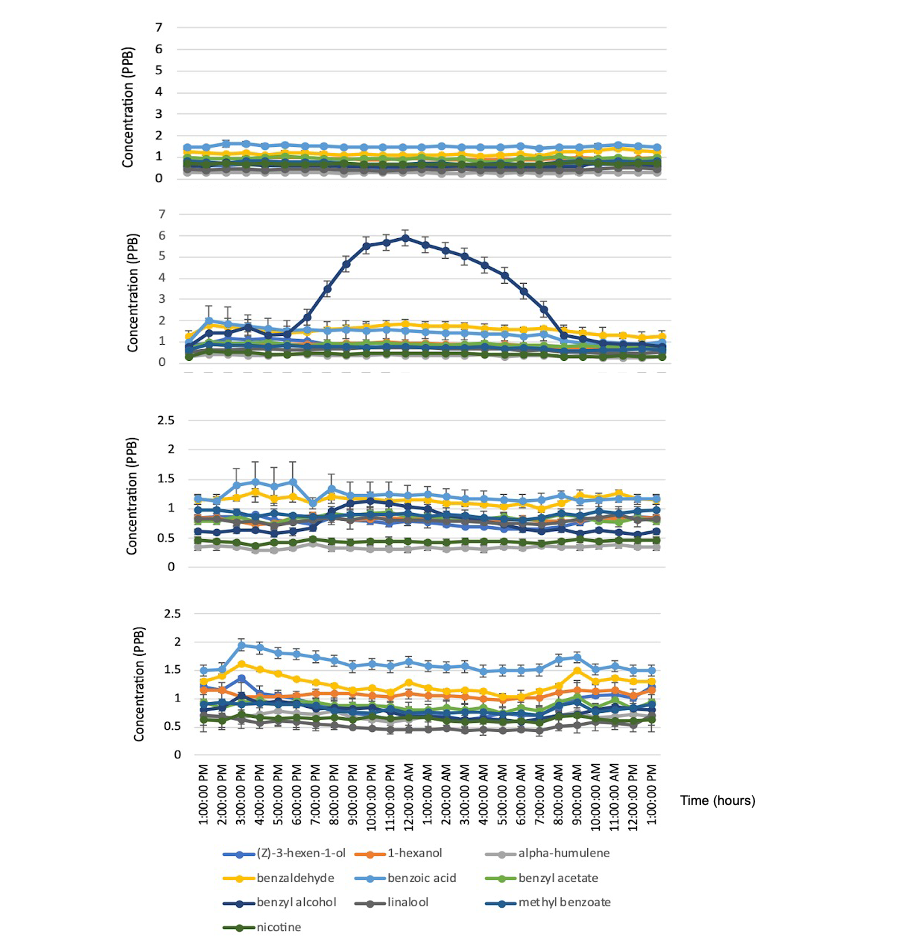

**Fig. S1.** Profiles of average emission of selected compounds in green leaf and floral headspace of LAB and QLD over a 24-hr period. (**A**) LAB floral headspace (**B**) QLD floral headspace (**C**) LAB green leaf headspace (**D**) QLD green leaf headspace.

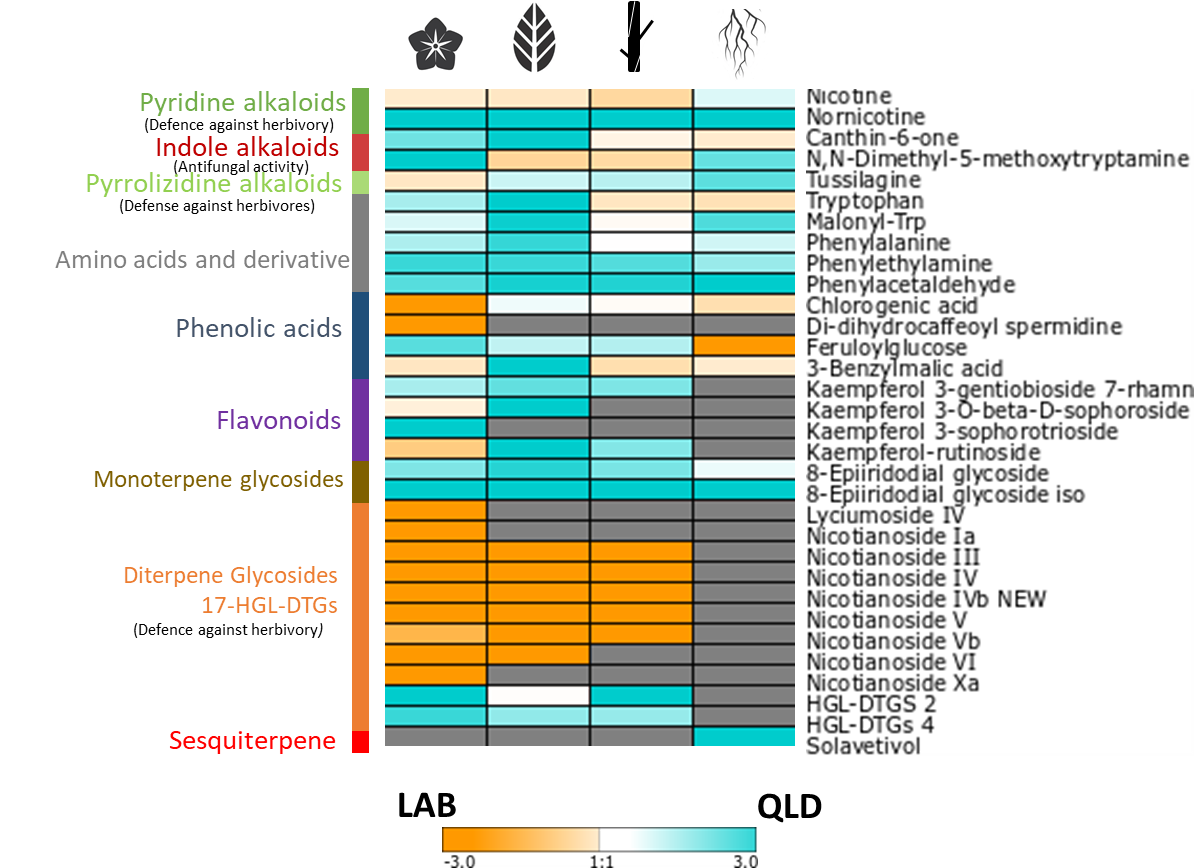

**Fig. S2.** Differentially accumulated metabolites in semi-polar extracts of *N. benthamiana* LAB vs QLD tissues analysed by liquid chromatography/high resolution mass spectrometry (LC/HESI/MS). Different colours indicate relative levels in LAB vs QLD, grey shaded areas not detectable levels.

**A**

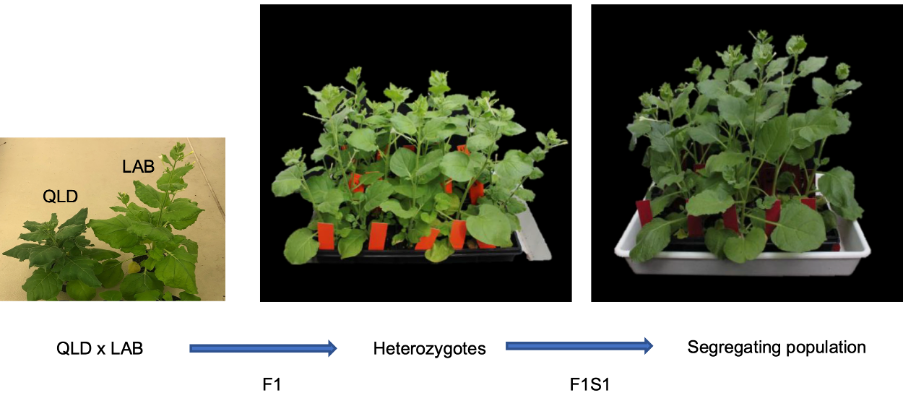

**B**

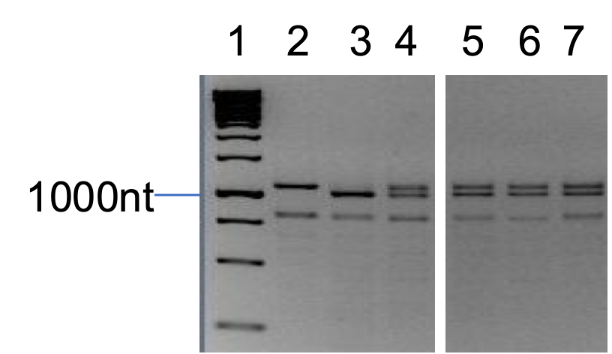

**Figure S3.** Inter-fertility of LAB and QLD. (**A**) Pollen from LAB was used to fertilise emasculated flowers of QLD and vice-versa. Both directions produced capsules containing >50 seeds. Seeds were germinated and grown in soil to set F1S1 seed or backcrossed with LAB or QLD. All crosses and selfing-derived seed gave healthy fertile plants with a variety of morphologies. (**B**) Hybridisation was confirmed by testing progeny of the initial LAB x QLD cross by PCR across the Rdr1 locus, which has a homozygous 72 bp insertion in the LAB background. Lanes 2,3 and 4 are LAB (homozygous insertion), QLD (homozygous no insertion), and a known heterozygote, respectively. Lanes 5, 6 and 7 are samples from three plants generated in the LAB x QLD cross.

**A**

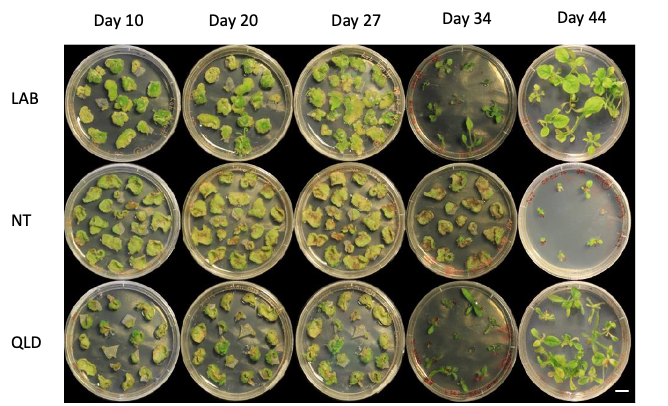

B

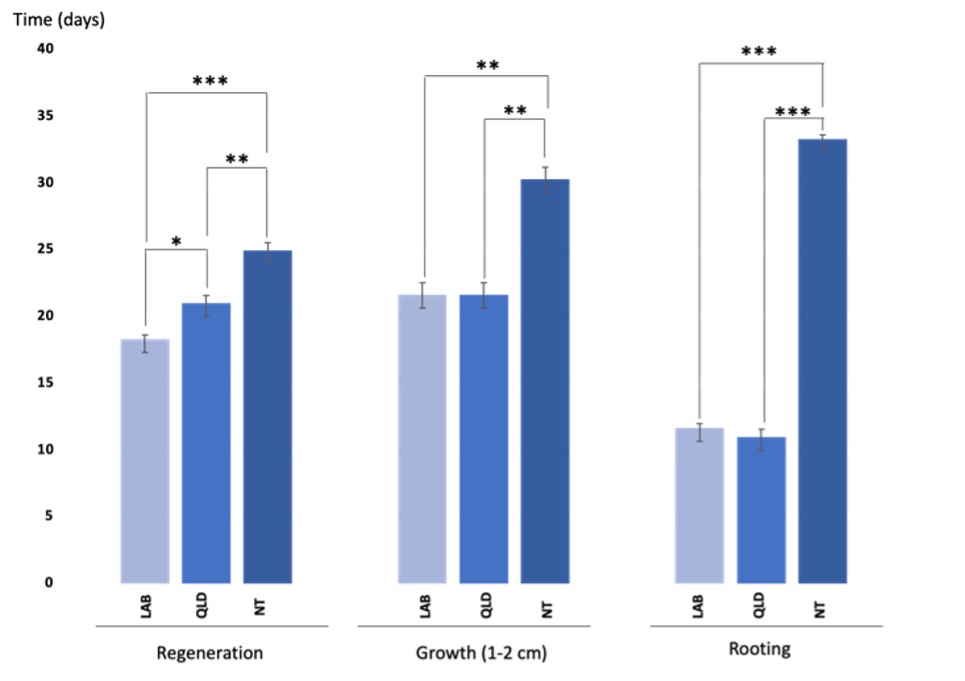

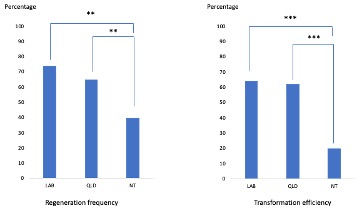

**Figure S4.** Comparison of transformation efficiencies between LAB, QLD and Northern Territory (NT) ecotypes. (**A)** Regeneration, selection, shoot development, and root development of LAB, NT and QLD ecotypes post-transformation with a 35S:Cas9 cassette and kanamycin selectable marker (scale bar represents 1 cm). The progression of transformation is indicated by the dates on top of the image. (**B**) Comparison of time taken for regeneration, growth (1-2 cm shoots) and rooting of LAB, QLD and NT (p value < 0.05*, p value < 0.01* *, p value < 0.001***). Three leaves per each biological replicate were infiltrated. Three replicates per each ecotype was used. (**C**) Comparison of regeneration frequency and transformation efficiency of LAB, QLD and NT (p value < 0.05*, p value < 0.01* *, p value < 0.001***). Independent positive transformants of LAB n=72, QLD n=74 and NT n=21(a single sister plant derived from a one single callus) were used to calculate the transformation efficiency. No significant differences were detected between LAB and QLD.

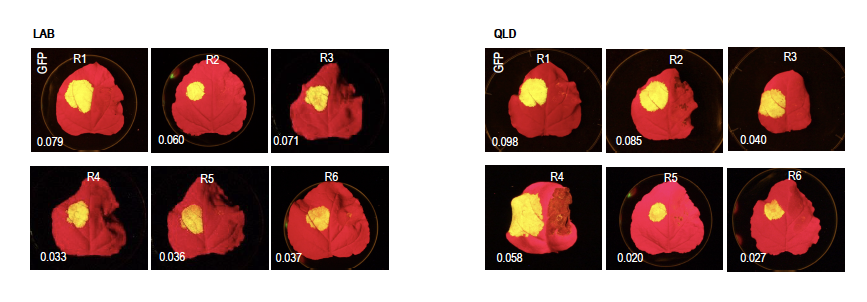

**Figure S5.** Comparison of transient expression of green fluorescence protein (GFP) in LAB and QLD by agroinfiltration of a 35S:GFP:HSP construct. GFP expression levels are represented underneath each leaf as log2(fold-changes). Chi-square and two sample equal variance T-test of LAB and QLD (6 replicates for each ecotype) gave p=0.99995 and p=0.89832, respectively, showing that there is no significant difference between GFP expression levels in the two ecotypes.

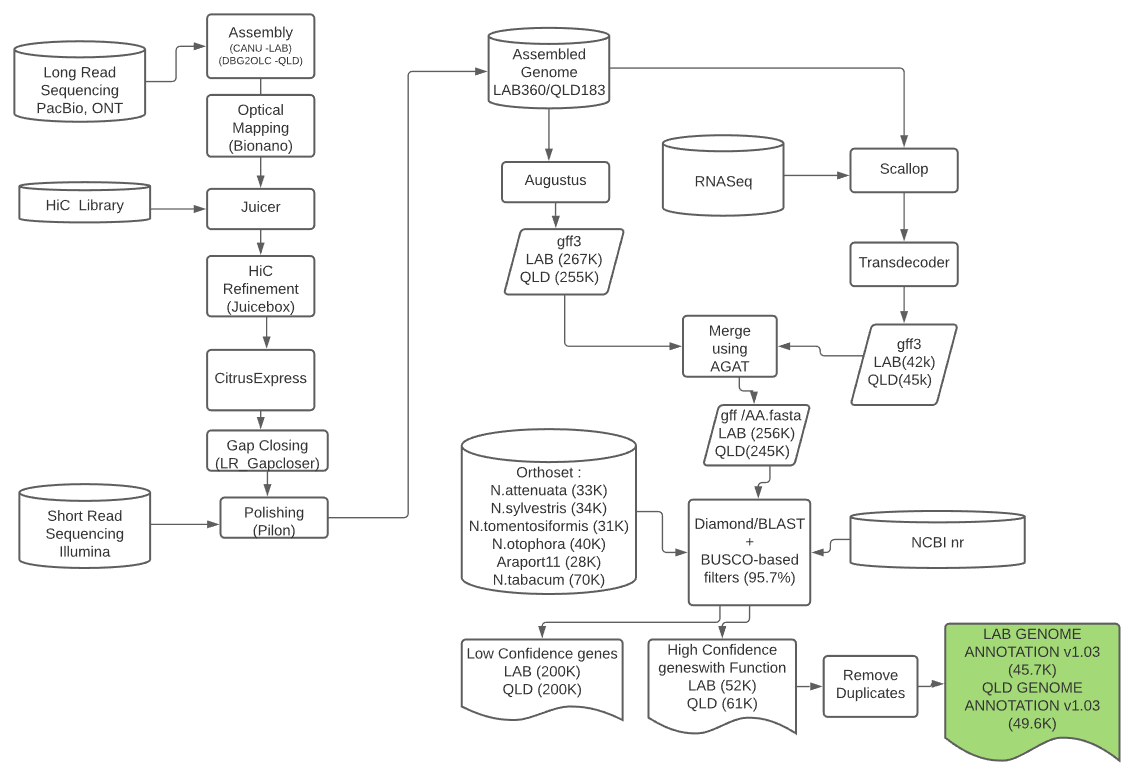

**Figure S6.** Assembly and annotation pipeline of the *N. benthamiana* genome based on PacBio, ONT long reads, bionano, and chromosome conformation capture (Hi-C). DNA was extracted from purified leaf nuclei (see online methods) for both LAB and QLD. Illumina sequencing was performed on a HiSeq2500, giving ~200 Gb of paired end 150bp reads. Long read sequencing was performed using both PacBio (Sequel and RSII platforms; ~160 Gb per genome) and ~20 Gb ONT (Oxford nanopore R9.4). For wild accessions, 50x coverage (~150Gb) was obtained using the Hiseq2500 platform.

A

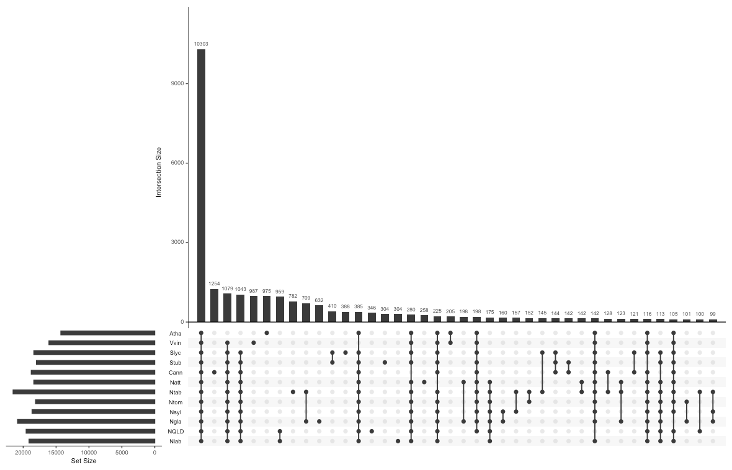

**Figure S7**  Upset plot showing orthologous groups between LAB, QLD, N.tabacum, N.sylvestris, N.tomentosiformis, N.glauca, A.thaliana, V.vinifera S.lycopersicum and S. tuberosum. Filled dots (black) denote the presence, and empty dots (grey) indicate the absence of orthologous groups in each specie. The plot was generated using UpSetR package in R(Conway et al. 2017).

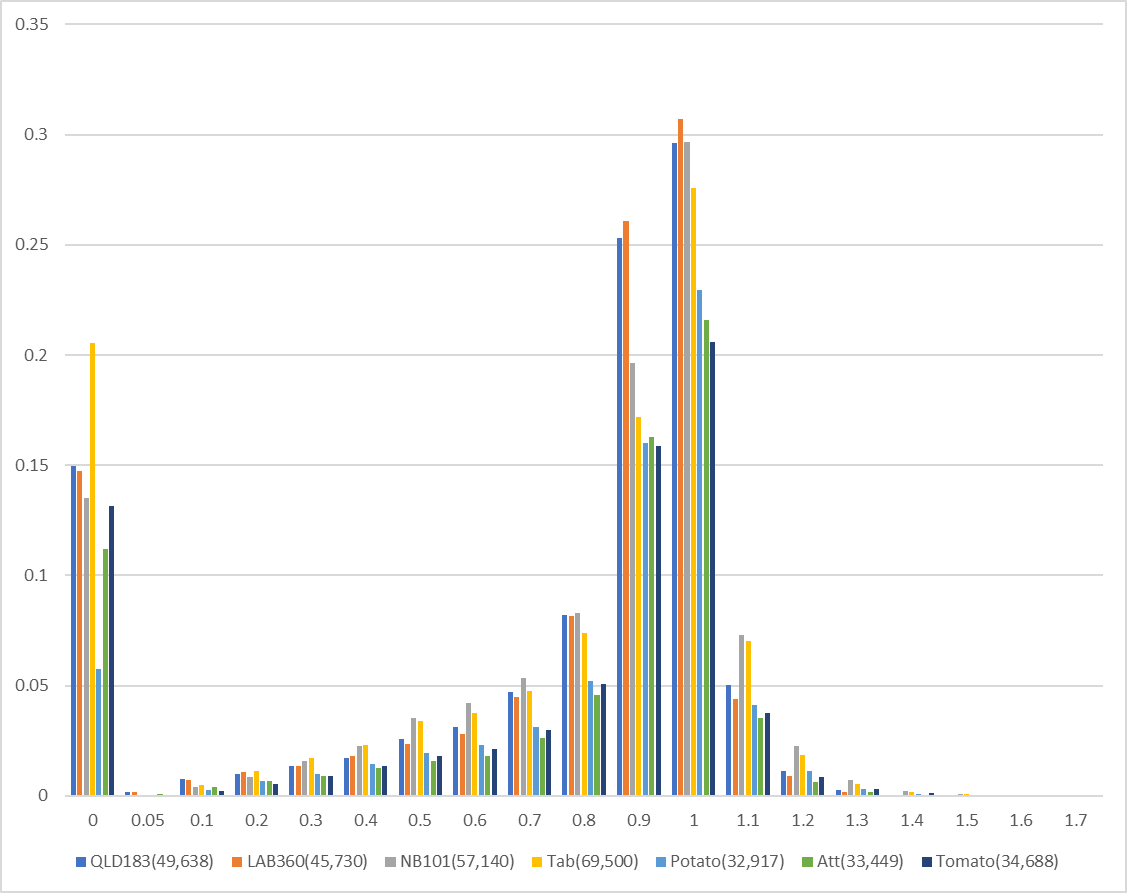

**Figure S8**. Completeness and quality of the LAB and QLD annotations. The predicted protein lengths of QLD, LAB, NB101, Tobacco, Potato, *N. attenuata* and Tomato were expressed as ratios compared to their Arabidopsis best hits (E-10). The 0 bin contains proteins that do not have an Arabidopsis match.

**A**
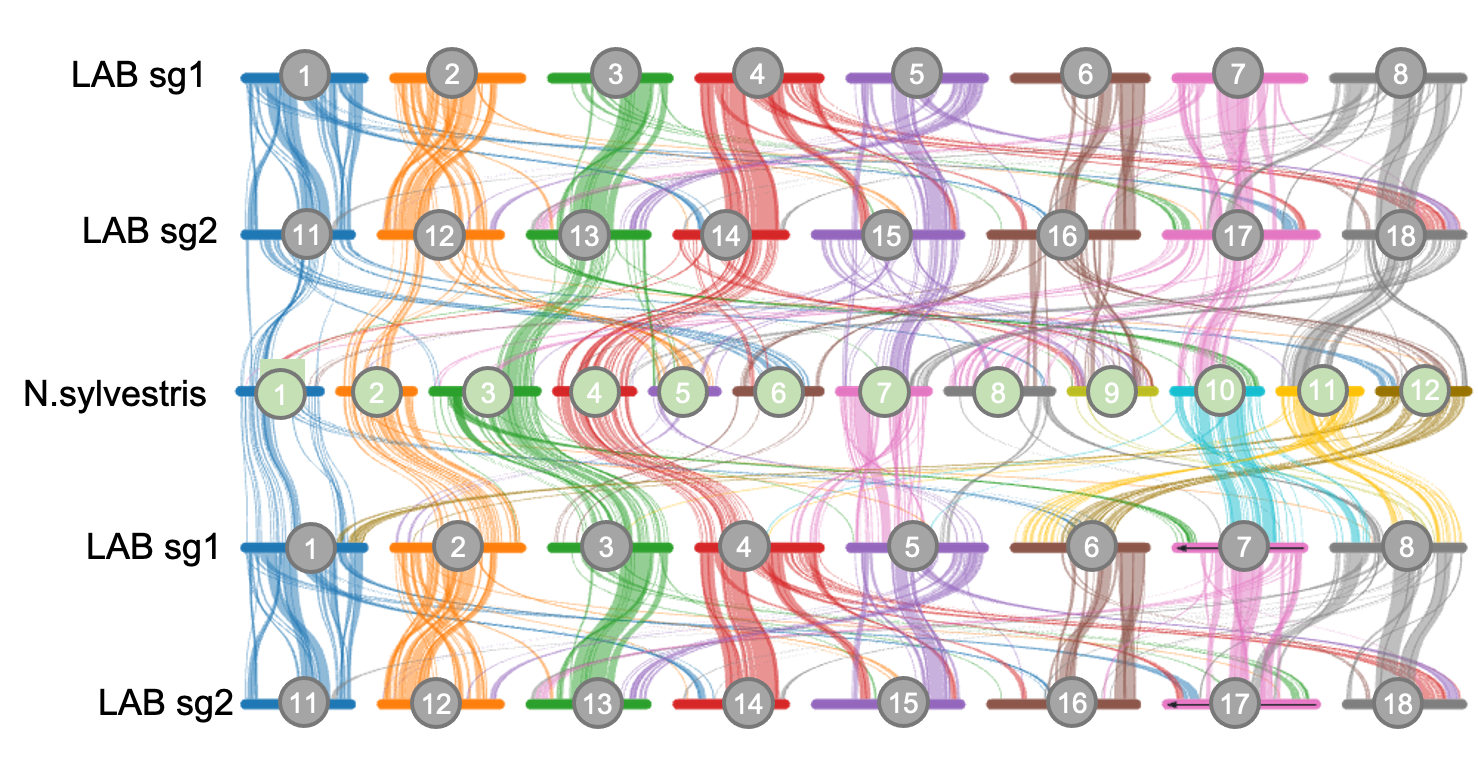

**B**

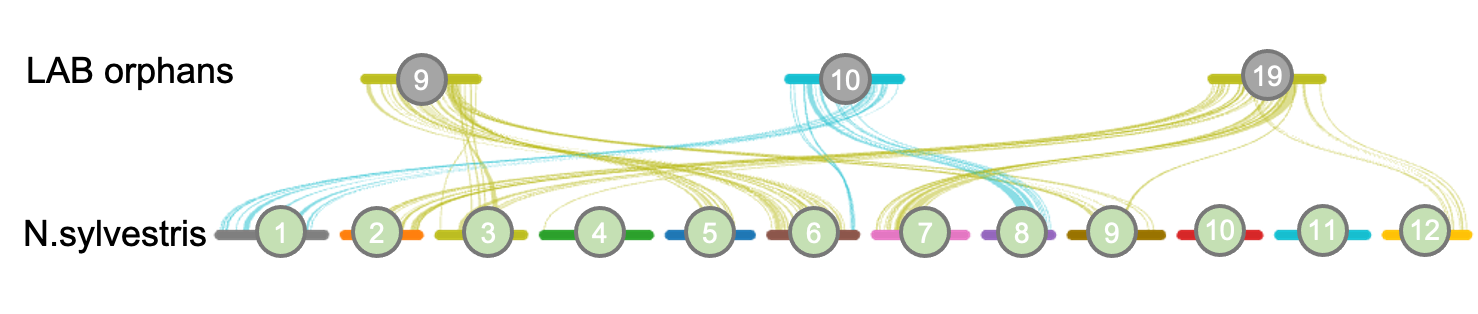

**Figure S9.** Syntenic blocks between the two subgenomes of *N. benthamiana*  and with the *N.sylvestris-*derived subgenome of *N.tabacum.*

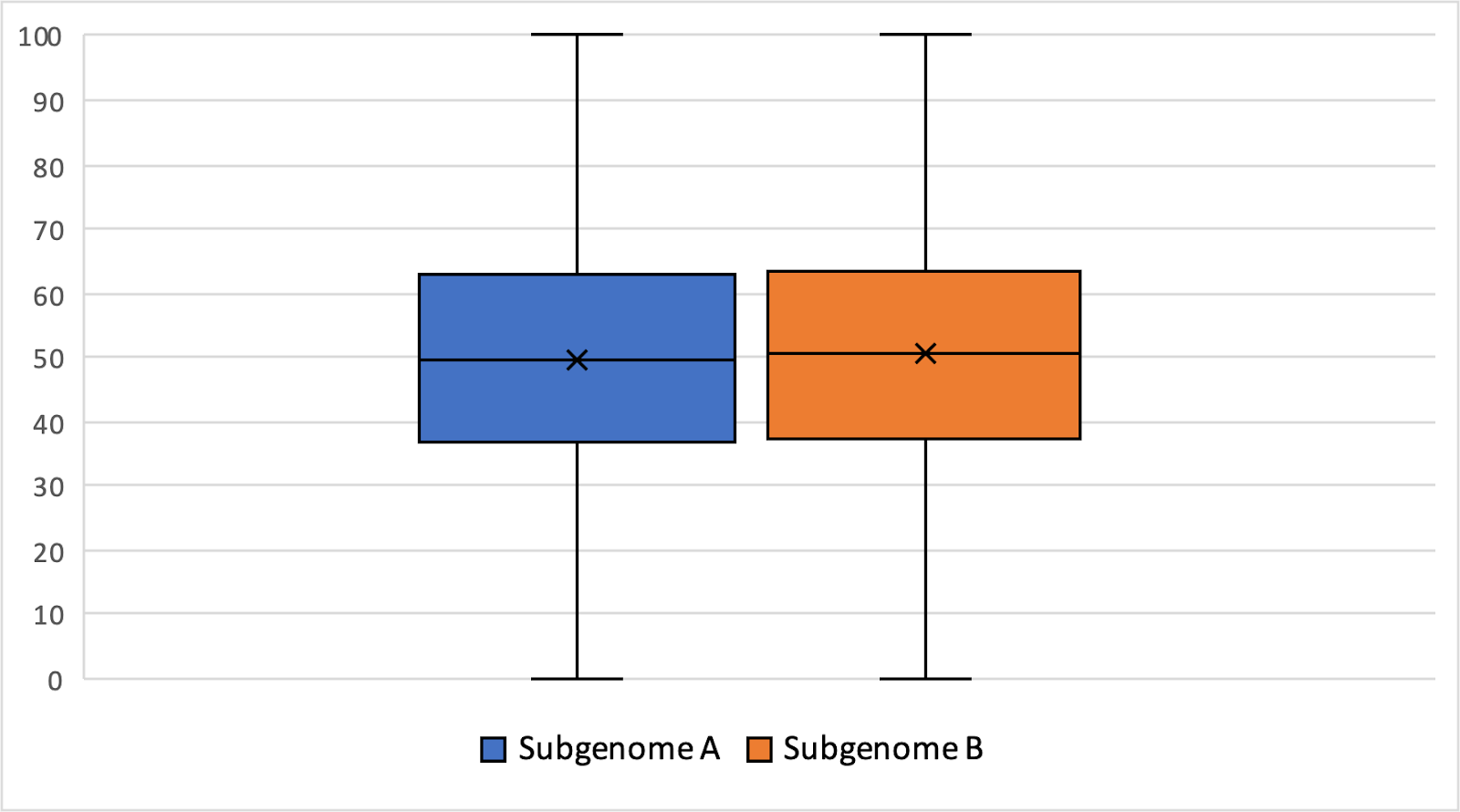

**Figure S10. Average relative homeolog expression in subgenomes of *N. benthamiana*.** No significant subgenome dominance (based on the homeolog expression bias) was observed (Kruskal-Wallis p value > 0.1).

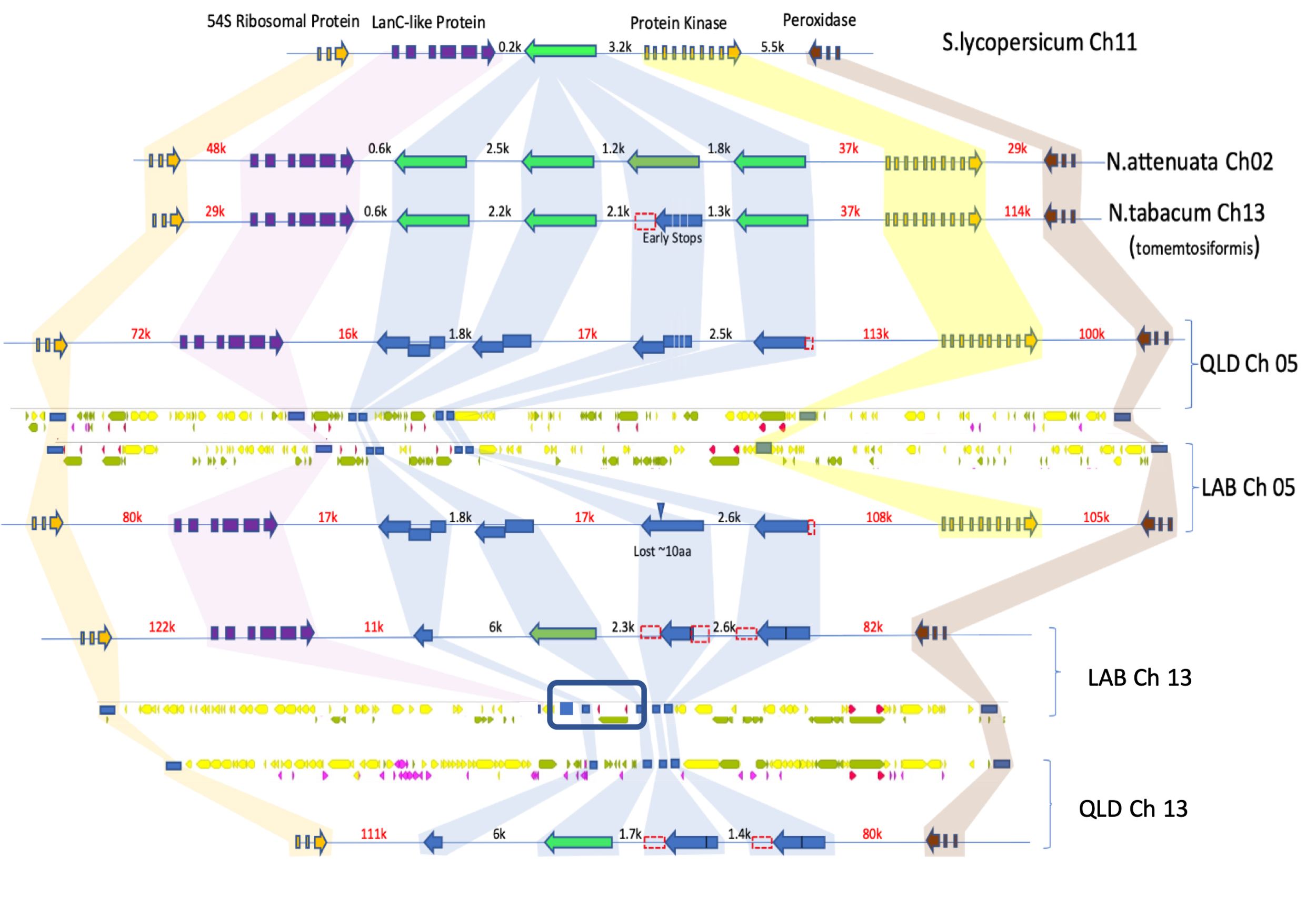

**Figure S11. Synteny of RPM1-like loci in tomato, *N. attenuata*, *N.tabacum*, LAB and QLD.** Gene arrangement in cartoon form representing RPM1-like genes (functional - bright green), possibly functional (dark green), pseudogenes (blue). Surrounding syntenic genes are shown in orange, purple, yellow and brown. Orthology/homeology relationships are indicated by colored shading. Distances between genes indicated (black text)< 15kbp; (redtext) >15kbp. Only LTR-transposable elements are shown. Yellow blocks represent GYPSY elements and green blocks represent COPIA elements. Red triangles represent LTR repeat regions that flank either a GYPSY or a COPIA element. These elements are likely to be nearly complete and can be considered possible autonomous elements. The rectangular red blocks flank unknown LTR-TE elements. Unknown TEs are elements that are recognisable as a LTR element but are not able to be classified as either a COPIA or GYPSY element due to irregularities in internal sequences for that element. These are likely to represent non-autonomous elements. Those elements not flanked by LTR sequences are highly fragmented non-functional elements. The blue rectangular boxes highlight the location of the genes annotated in the tracks above and below the TE annotation tracks.

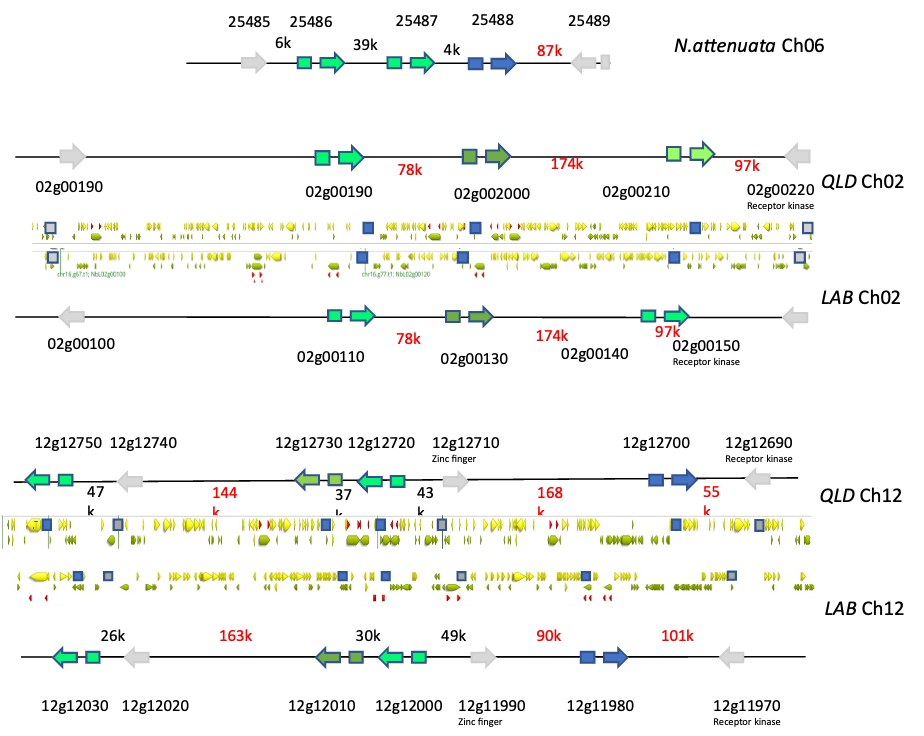

**Figure S12. CYP82E Nicotine demethylase loci in N.attenuata, LAB and QLD.** Gene arrangement in cartoon form representing CYP82E-like genes (functional - bright green), possibly functional (dark green), defective (blue). Distances between genes indicated (black text)< 15kbp; (redtext) >15kbp.

TE annotation tracks for LAB and QLD were prepared using annotation data from the EDTA TE annotation pipeline (see online methods) and Geneious Prime software (Geneious Prime® 2023.0.1; https://www.geneious.com). Only LTR-transposable elements are shown. Yellow blocks represent GYPSY elements and green blocks represent COPIA elements. The size of each block is proportional to the number of base-pairs annotated for that element. Red triangles represent LTR repeat regions that flank either a GYPSY or COPIA element. These elements are likely to be nearly complete and can be considered possible autonomous elements. The rectangular red blocks flank unknown LTR-TE elements. Unknown TEs are elements that are recognisable as a LTR element but are not able to be classified as either a COPIA or GYPSY element due to irregularities in internal sequences for that element. These are likely to represent non-autonomous elements. Those elements not flanked by LTR sequences are highly fragmented non-functional elements. The blue rectangular boxes highlight the location of the genes annotated in the tracks above and below the TE annotation tracks.

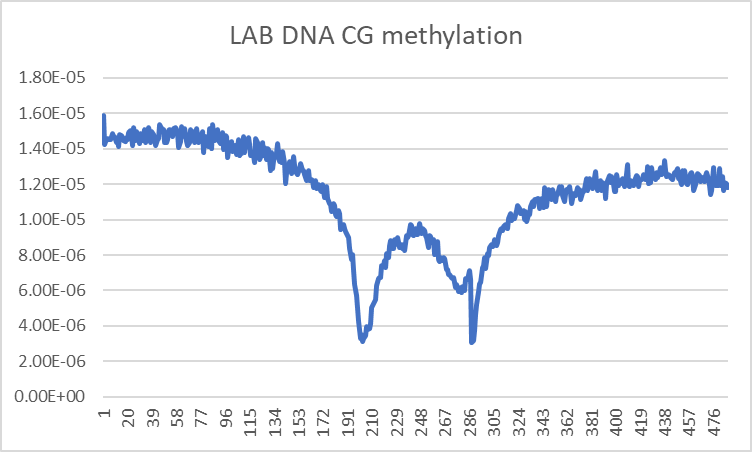

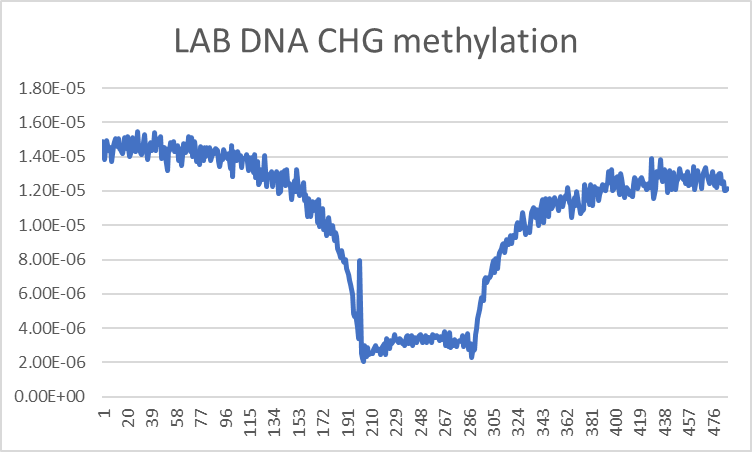

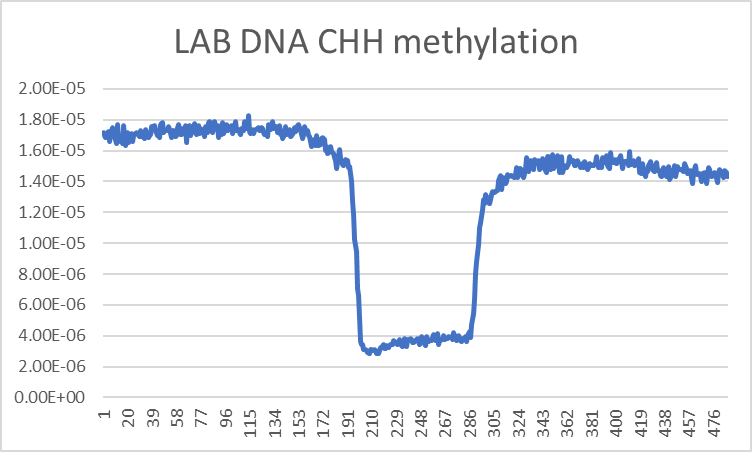

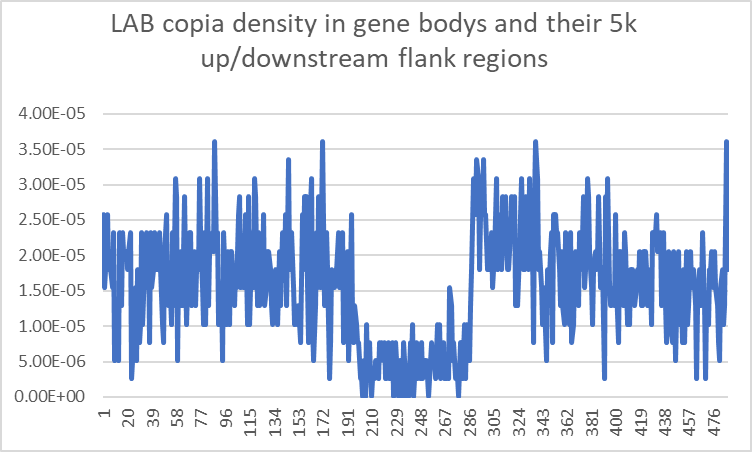

**Figure S13. Copia element insertions and methylation profiles in the proximity of LAB genes.** Vertical axis shows the fraction of Copia elements per bin of size 25bp. The horizontal axis is separated into three segments: (1): from 1 to 200(=5000bp/25bp) upstream flanking region. (2) from 200 to 300 (average gene length 2517bp/25bp): coding region. (3) from 300 to 500(=5000+5000+2517/25) downstream flanking region. Different gene lengths were normalised to fit the average gene length.
